# HEXIM1/P-TEFb complex controls RNA polymerase II pause release and immediate early gene induction following neuronal depolarization

**DOI:** 10.1101/2024.09.27.615234

**Authors:** Myo Htet, Camila Estay-Olmos, Lan Hu, Yiyang Wu, Brian E. Powers, Clorissa D. Campbell, Aishwarya Rameshwar, Mohamed R. Ahmed, Timothy J. Hohman, Yanling Wang, Julie A. Schneider, David A. Bennett, Vilas Menon, Philip L. De Jager, Garrett A. Kaas, Roger J. Colbran, Celeste B. Greer

## Abstract

Cognitive processes require *de novo* gene transcription in neurons. Memory requires the rapid and robust transcription of a class of genes called immediate early genes (IEGs). IEG transcription is facilitated by the formation of a poised basal state, in which RNA polymerase II (RNAP2) initiates transcription, but remains paused downstream of the promoter. Upon neuronal depolarization, the paused RNAP2 is released to complete the synthesis of messenger RNA (mRNA) transcripts, a process stimulated by positive transcription elongation factor b (P-TEFb). In many cell types, P-TEFb is sequestered into a large inactive complex containing Hexamethylene bisacetamide inducible 1 (HEXIM1), but the impact of this interaction on neuronal gene transcription is not yet fully understood. In this study, we found that neuronal expression levels of *HEXIM1* mRNA are highly correlated with impaired cognition in Alzheimer’s disease. It is also induced in the hippocampus during memory formation, and following depolarization in neurons. The role of HEXIM1 in neuronal gene transcription was then explored in murine neuronal cultures where we found that calcium frees P-TEFb from the HEXIM1 inhibitory complex. Modulation of P-TEFb by inhibiting the activity of the cyclin-dependent kinase 9 (CDK9) subunit of this complex significantly impacts IEG induction, particularly during repeated depolarization. Our findings indicate that HEXIM1 in complex with P-TEFb plays an important role in establishing and resetting the poised RNAP2 state, enabling efficient activation of genes necessary for synaptic plasticity.

## Introduction

Learning and memory formation require carefully orchestrated changes in gene transcription programs carried out by RNA polymerase II (RNAP2) that are stimulated by increased synaptic activity and calcium influx (1–4). Transcription of a set of immediate early genes (IEGs) is activated in the period immediately following neuronal depolarization, producing messenger RNAs (mRNAs) that encode a variety of protein types. Two major categories are genes that encourage neuronal plasticity such as (Activity regulated cytoskeleton associated protein, *Arc*; *Homer1*; etc.) and transcription factors such as (*Fos*; Early growth response factor 1, *Egr1*; *Jun*; Neuronal PAS domain protein 4, *Npas4*; Nuclear receptor subfamily 4 group A member 2, *Nr4a2*; etc.) that promote the transcription of additional genes. IEGs are induced in the hippocampus during a variety of cognitive processes, including fear conditioning (5–9). Recent work has identified that repeated neuronal stimulation leads to damped successive transcriptional responses of IEGs (10). However, molecular mechanisms underlying the dampened response to serial stimuli remain unknown. Moreover, the potential recovery from transcriptional dampening and the role of transcriptional dampening in human disease has not been investigated.

RNAP2-dependent synthesis of new mRNAs occurs in phases as the polymerase transitions between initiation, pausing, elongation, and termination. IEGs are paused in unstimulated neurons, and elongation of RNAP2 is elicited by depolarizing stimuli (11). Positive transcription elongation factor b (P-TEFb) releases paused RNAP2 into the elongation phase (12). This is accomplished in part through its direct phosphorylation of its largest subunit at the second serine in the heptad repeat of its C-terminal tail (RNAP2-pS2) (13). P-TEFb is a heterodimer of cyclin-dependent kinase 9 (CDK9) and either Cyclin T1 or T2 (CCNT1 or CCNT2). When actively inducing elongation, P-TEFb associates with bromodomain-containing protein 4 (BRD4) (14), an acetyl-lysine binding protein that targets active P-TEFb to acetylated histones surrounding active gene promoters that contain paused RNAP2. Alternatively, association of the P-TEFb heterodimer with an inhibitory complex containing Hexamethylene bisacetamide inducible 1 (HEXIM1) restricts P-TEFb activity (15). This transcriptional silencing complex also contains methylphosphate capping enzyme (MEPCE), La-related protein 7 (LARP7), and the non-coding RNA RN7SK (16) (**Fig.1A**). Release of P-TEFb from the silencing effects of the HEXIM1 containing complex regulates inducible gene expression in response to stimuli like ultraviolet light exposure or virus infection in cancerous and other non-neuronal cell (17–19). Despite the importance of this complex in transcriptional regulation in other cell types, the specific role of the HEXIM1 in the brain is only recently beginning to be investigated (20), and the effect of HEXIM1 on the important stimulus-dependent expression of IEGs in neurons is previously unexplored.

**Figure 1.**
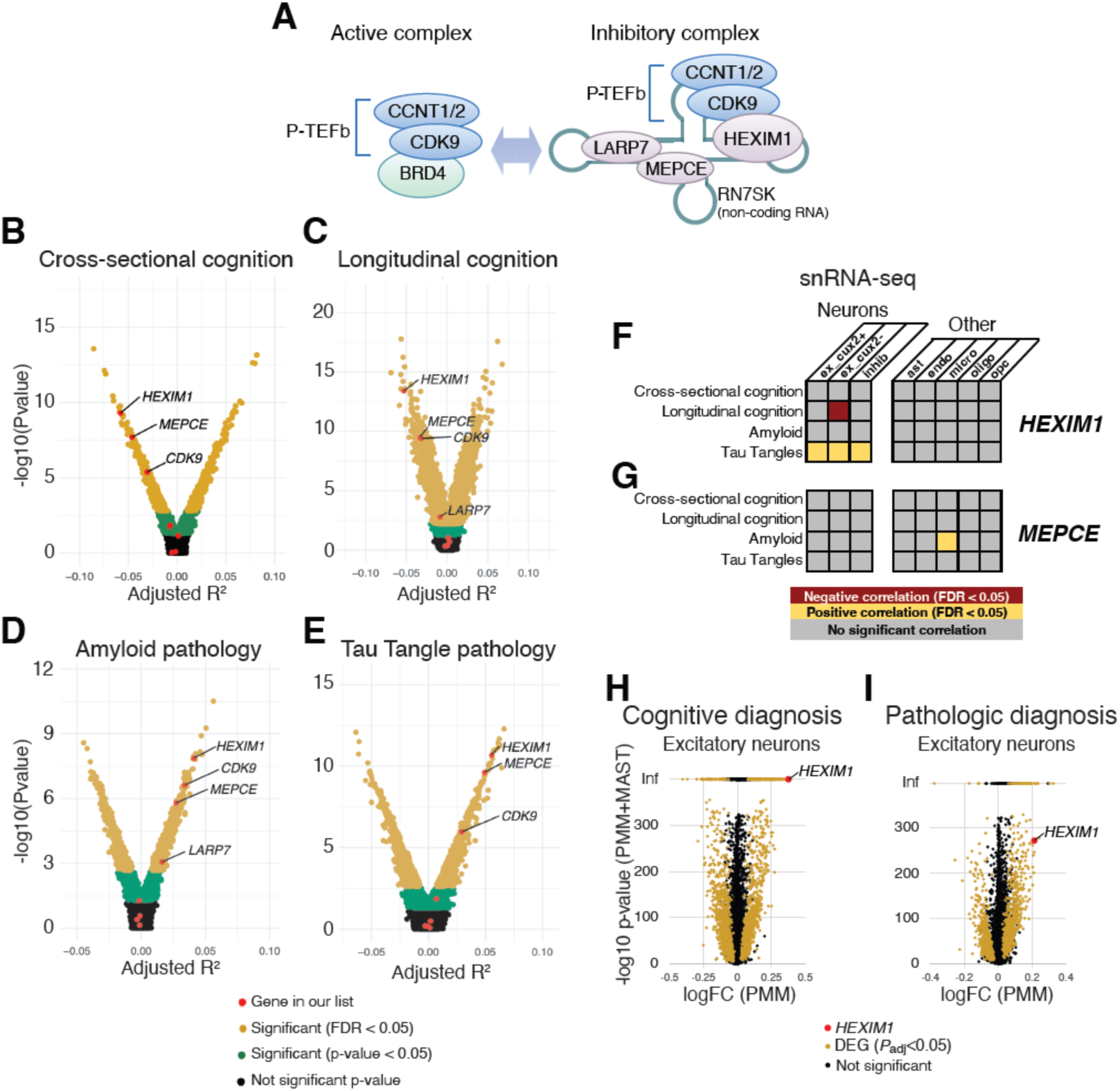
P-TEFb regulatory complex components and their correlations with characteristics of AD in bulk RNA-seq and snRNA-seq. (**A**) Cartoon depicting components of the active and inhibitory P-TEFb complexes. (**B-E**) Correlations (R^2^) between expression of *BRD4*, *CCNT1*, *CCNT2*, *CDK9*, *HEXIM1*, *LARP7*, and *MEPCE* mRNAs and *RN7SK* non-coding RNA with (**B**) cross-sectional cognition, (**C**) longitudinal cognition, (**D**) amyloid pathology, and (**E**) tau tangle pathology in bulk tissue RNA-seq from the head of the caudate nucleus. Individual dots represent a single gene, and genes in the yellow region of the volcano plots are significantly correlated with the indicated pathology (FDR < 0.05). (**F-G**) snRNA-seq correlations with AD pathologies in different cell types in DLPFC. Significant correlations of (**F**) *HEXIM1* and (**G**) *MEPCE* mRNAs are depicted in yellow (positive correlation) or red (negative correlation). ex = excitatory neurons, inhib = inhibitory neurons, ast = astrocytes, endo = endothelial cells, micro = microglia, oligo = oligodendrocytes, opc = oligodendrocyte progenitor cells. (**H-I**) Correlation between *HEXIM1* expression and (**H**) cognitive diagnosis of AD and (**I**) pathological diagnosis of AD in excitatory neurons across six brain regions analyzed by snRNA-seq in an independent cohort as data reported in **F** and **G**. Log fold change (FC) is plotted against the -log10 of a summed p-value for two independent statistical analyses of the data (see Experimental Procedures). Yellow dots represent differentially expressed genes (DEG), which were defined as P_adj_<0.05 in both analyses, with the directionality of the change being concordant. Inf = infinity. *HEXIM1* was identified as a DEG in these datasets and is highlighted in red.

Alzheimer’s disease (AD) is an aging-associated neurodegenerative disorder that leads to progressive cognitive impairment and memory loss. The accumulation of amyloid beta (Aβ) plaques and neurofibrillary tangles comprised of Tau protein aggregates in the brain are well-established biomarkers of this destructive disease (21). There is a growing appreciation that AD is also associated with alterations in gene expression in a variety of cell types in the brain, and transcription is especially perturbed in neurons (22). Neuronal IEGs are misregulated in aging and AD (23–28), and IEG expression in AD rodent models can be rescued by modulating the level of RNAP2 pausing (29, 30). Therefore, factors regulating pausing could be effective AD targets, and there are several examples of pause release inhibitors improving cognitive functions in AD model organisms. First, flavopiridol, a P-TEFb inhibitor, rescued passive avoidance and object recognition memory deficits in mice induced by the injection of Aβ oligomers (31). BRD4 inhibition has also been shown to improve memory in some (30, 32–34), but not all (35) rodent models of Aβ accumulation. Moreover, inhibitors of histone deacetylases (HDACs), which decrease transcription pause release (36–39), improve memory in mice with Aβ accumulation (40–44). These prior observations led us to hypothesize that release of paused RNAP2 is too high in AD, and that these effective treatments corrected the impairment in this regulatory step. Because drugs that promote transcription elongation also sometimes induce compensatory increases in *HEXIM1* transcription (45), we further hypothesized that dysregulation of RNAP2 pause-release in AD could be due to alterations in the transcription of P-TEFb regulators.

We set out to first determine whether P-TEFb and its regulators are altered in AD. Using postmortem human data (46), we found that the expression levels of certain subunits of P-TEFb and components of the P-TEFb inhibitory complex are correlated with AD pathology across several brain regions. In particular, *HEXIM1* is highly dysregulated, and single nuclei RNA sequencing (snRNA-seq) revealed correlations between its neuronal expression levels and both declining cognition and increased pathology. Therefore, we investigated the contributions of HEXIM1 and P-TEFb to neuronal gene transcription using primary mouse hippocampal neuron cultures. Using pharmacological and genetic approaches, we demonstrate that HEXIM1 sequestration of P-TEFb at IEG promoters primes neurons for activity-dependent transcription, and calcium dissociates this complex. After IEG activation, there is a period where neurons cannot reactivate certain IEGs as robustly, but they eventually recover after several hours. This timeline coincides with a period of decreased HEXIM1 protein expression, and the dampened response of IEGs to stimuli during this period is P-TEFb-dependent. Overall, our work shows that expression of *HEXIM1* is dysregulated in AD and implicates HEXIM1 as an important modulator of the gene expression changes that follow neuronal depolarizations.

## Results

### Neuronal HEXIM1 mRNA expression correlates with Alzheimer’s disease pathology

In light of the possible link between AD and regulators of transcriptional pausing, we sought to understand how cognition and AD-associated pathologies correlated with the expression of mRNAs encoding some known regulators of RNAP2 pause release in the AD brain. Subjects enrolled in the Religious Orders Study/Memory and Aging Project (ROS/MAP) study provided antemortem cognitive assessments. Postmortem, their brain tissue was analyzed for pathological markers and were subjected to transcriptional analysis by bulk tissue RNA-seq and single nuclei RNA-seq (46). Within this published dataset, we analyzed correlations between mRNA expression levels of *BRD4*, *CCNT1*, *CCNT2*, *CDK9*, *HEXIM1*, *LARP7*, *MEPCE*, and *RN7SK*, which represent several P-TEFb components and regulators, with the patient’s AD-associated cognitive changes and neuropathology severity in the head of the caudate nucleus (CN). Severity of the neuropathology of amyloid and tau-tangles was determined across eight forebrain regions using immunohistochemistry (47). We found that cognitive scores in study participants from their final clinical evaluation prior to death (Cross-sectional cognition; **Fig. 1B**), changes in cognitive assessment scores across time (Longitudinal cognition; **Fig. 1C**), amyloid pathology (**Fig. 1D**) and tau tangle pathology (**Fig. 1E**) were significantly correlated with several of these genes and AD-pathologies (FDR < 0.05). Among the data from the CN, *HEXIM1* provided the strongest correlations with all these pathological measures we examined: a significant negative correlation with cognitive scores (meaning increased expression is associated with worse cognitive performance) and a significant positive correlation with amyloid and Tau pathologies (meaning increased expression is associated with more pathology). *MEPCE*, another of the inhibitory complex components, and *CDK9*, encoding the kinase subunit of P-TEFb, also significantly correlated in the same directions as *HEXIM1* in all four measures (**Fig. 1B-E**). *LARP7* correlated with poorer longitudinal cognition and increased amyloid pathology (**Fig. 1C-D**). Components of the inhibitory complex (including subunits of P-TEFb) correlated with AD pathology in two other brain areas (dorsolateral prefrontal cortex (DLPFC), and posterior cingulate cortex (PCC)) (**Supp. Fig. 1A-H**). Of note, *HEXIM1* and *MEPCE* expression levels significantly correlated with worse cognition and AD pathology in every brain area examined (summarized in **Supp. Fig. 1I**). *HEXIM2*, a gene paralog that can sometimes compensate for *HEXIM1* (48), does not correlate with AD pathology (**Supp. Fig. 1J**). We also examined other interactors of CDK9, and though some correlations were found (**Supp Fig. 1 K-L**), none were as consistently correlated with AD as *HEXIM1* and *MEPCE* across brain regions and pathologies.

The correlations discussed so far were identified from transcriptional analyses of bulk tissue, representing a mix of cell types. Therefore, we sought to investigate the cell type specificity of these correlations, as well as their reproducibility in a separate data set generated from some of the same samples. To this end, we analyzed expression levels of the hits we had the highest confidence in from the bulk sequencing data, *HEXIM1* and *MEPCE,* in snRNA-seq data sets generated from the ROS/MAP samples. Neurons were clustered into three types (excitatory cut-like homeobox 2 (cux2) positive (ex cux2+), excitatory cux2 negative (ex cux2-), and inhibitory (inhib)). Cux2 is expressed in layers II-IV of the cortex (49, 50), and can be used as a marker to divide large datasets of excitatory neurons into populations representing upper and lower cortical layers (51). Additionally, five non-neuronal cell types (astrocytes (ast), endothelial cells (endo), microglia (micro), oligodendrocytes (oligo), and oligodendrocyte progenitor cells (opc)) were defined. Amongst the components of the P-TEFb inhibitory complex, *HEXIM1* levels in multiple neuronal subtypes correlated with several aspects of AD pathology (**Fig. 1F**). In particular, the longitudinal change in cognition was negatively correlated with *HEXIM1* expression levels in ex cux2-(deeper layer) neurons, and tangles were positively correlated with *HEXIM1* expression in all neuronal subtypes (**Fig. 1F**). *MEPCE* expression is positively correlated with amyloid pathology in micro cells (**Fig. 1G**) in this dataset.

snRNA-seq data set was recently generated from another cohort of control and AD-affected individuals (52). In this study, excitatory neurons were analyzed in a combined analysis of samples from six brain areas (entorhinal cortex, hippocampus, anterior thalamus, angular gyrus, midtemporal cortex, and prefrontal cortex) and differentially expressed genes (DEG) were identified. While *MEPCE* was not detected in excitatory neurons and therefore could not be analyzed, *HEXIM1* was correlated with cognitive (**Fig. 1H**), and pathological diagnoses of AD by National Institute on Aging (NIA)-Reagan diagnostic criteria (**Fig. 1I**). Taken together, multiple transcriptional analyses of two independent cohorts of AD samples suggest that expression levels of various components of the P-TEFb complex, and especially HEXIM1, are correlated with various aspects of AD pathology and/or symptoms.

### Hexim1 is an activity-dependent gene expressed in neuronal nuclei

Because of the correlations between *HEXIM1* mRNA and cognitive function in AD, we investigated whether its expression levels were also affected by learning and memory in a non-diseased animal model. We examined a publicly available RNA-seq dataset (53) generated from mice undergoing memory formation. We observed a significant increase in *Hexim1* mRNA expression in mouse hippocampal CA1 region 1 hour following a memory-inducing mild foot shock (**Fig. 2A**). IEGs including *Fos*, *Arc*, *Egr1* were also significantly induced (**Fig. 2B-D**). *Nr4a2* is trending up but was not significantly increased by our analysis (**Fig. 2E**). To directly determine whether HEXIM1 protein is expressed in neurons, we cultured primary hippocampal neurons from mice. HEXIM1 protein is co-expressed in cells stained by NeuN neuronal marker (**Fig. 2F**), and is mostly, but not exclusively, nuclear. To test if *Hexim1* mRNA expression is activity-dependent, as suggested by the fear conditioning experiment, we depolarized these cells with potassium chloride (KCl) (54), and saw a significant, but modest, increase in *Hexim1* mRNA (**Fig. 2G**). In these same samples, *Fos*, *Arc*, *Egr1*, and *Nr4a2* were significantly upregulated as well (**Fig. 2H-K**). To examine this transcriptional change in another cell system, we differentiated neuroblastoma Neuro2a cells (differentiated N2a; or dN2a) and likewise saw an increase in *Hexim1, Fos*, and *Egr1* mRNA in response to KCl (**Supp. Fig. 2**). These data suggests that neurons express HEXIM1 protein and dynamically regulate *Hexim1* mRNA expression in response to memory formation and neuronal depolarization.

**Figure 2.**
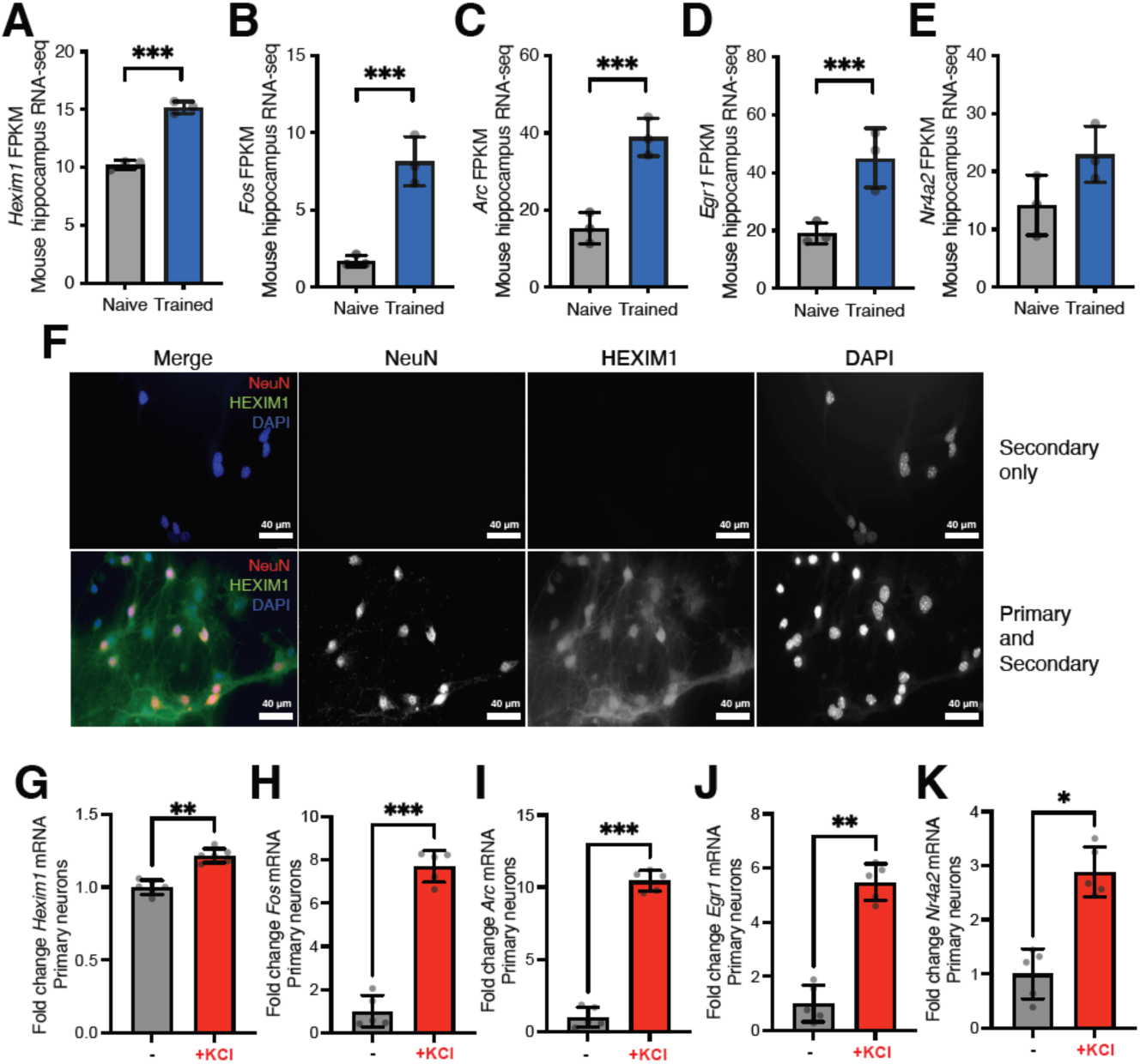
HEXIM1 is an activity-dependent gene that is expressed in hippocampal neurons. (**A-E**) Gene expression changes in mouse CA1 region of hippocampus 1 hr after mild foot shock. Average Fragments Per Kilobase of transcript per Million mapped reads (FPKM) aligning to (**A**) *Hexim1,* (**B**) *Fos,* (**C**) *Arc,* (**D**) *Egr1,* and (**E**) *Nr4a2* are depicted. Fisher’s exact test with FDR correction. \*\*\**FDR*<0.001. n=3 mice. (**F**) ICC of HEXIM1 in mouse primary neuron culture (hippocampal). Scale bar = 40µm. Top row depicts secondary antibody only control images, and bottom row depits fluorescent images of cells that had primary and secondary antibodies applied. Individual fluorescent channels are depicted in grayscale. In the merged image, NeuN is depected as red, HEXIM1 as green, and DAPI as blue. (**G-K**) RT-PCR of mRNA expression following 2 hr KCl stimulation in primary hippocampal neurons. Average expression of (**G**) *Hexim1,* (**H**) *Fos,* (**I**) *Arc,* (**J**) *Egr1,* and (**K**) *Nr4a2* mRNA all relative to the *Hprt* housekeeping gene are depicted. n=5 biological replicates. Paired two-tailed t tests. \**p*<0.05, \*\**p*<0.01, \*\*\**p*<0.001. Error bars represent standard deviation.

### HEXIM1 regulates stimulus-dependent IEG induction in neurons

We hypothesized that HEXIM1 might regulate IEGs in neurons because of its known role in regulating inducible gene expression in other cell types. To test this, we generated an adeno-associated virus of serotype 9 (AAV9) to overexpress Flag-tagged recombinant mouse *Hexim1* (F:*Hexim1*) under the control of the neuron-specific promoter human synapsin 1 (hSyn1). The recombinant protein runs at a slightly higher molecular weight due to the Flag tag (**Fig. 3A**). Both control virus and hSyn1:F:*Hexim1* expression viruses also express enhanced yellow fluorescence protein (eYFP) from a constitutive eukaryotic promoter, and viral transduction led to similar levels of eYFP expression **(Fig. 3B)**. HEXIM1 overexpression significantly dampened KCl-dependent IEG induction for *Fos* (**Fig. 3C**), *Arc* (**Fig. 3D**), *Egr1* (**Fig. 3E**), and *Nr4a2* (**Fig. 3F**).

**Figure 3.**
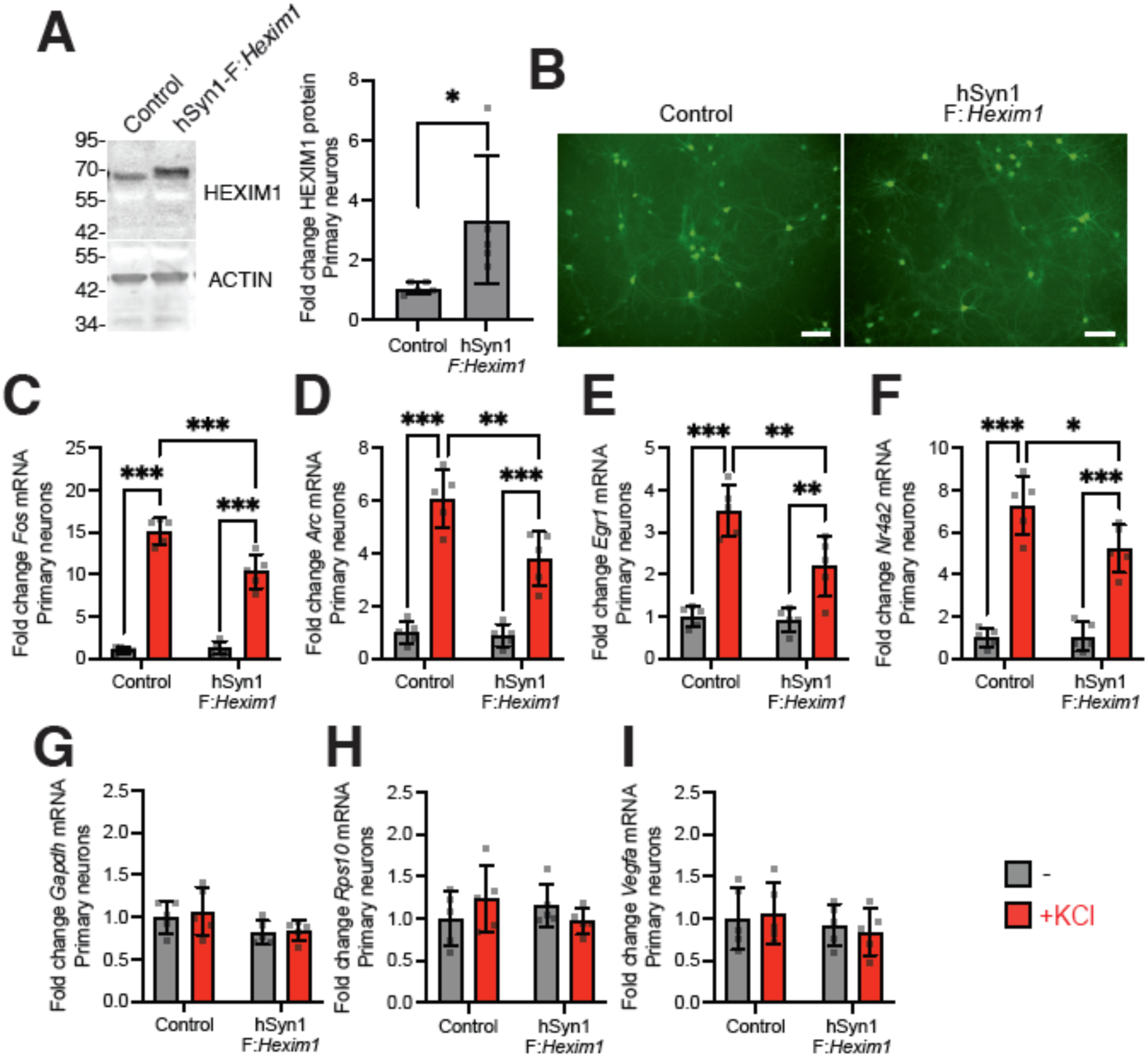
Effect of HEXIM1 overexpression on IEG induction in hippocampal primary neurons. (**A**) Representative Western blot of HEXIM1 and ACTIN in whole cell lysates following Control or hSyn1-FLAG:*Hexim1* AAV9 treatment and quantitation of HEXIM1 expression changes relative to ACTIN. Paired one-tailed t test. (**B**) eYFP expression following infection with Control or hSyn1-FLAG:*Hexim1* AAV9. Scale bar = 100µm. (**C-I**) mRNA levels (RT-PCR) after 1-week virus treatment followed by 2 hr KCl stimulation in primary hippocampal neurons. We analyzed (**C**) *Fos* (**D**) *Arc*, (**E**) *Egr1*, (**F**) *Nr4a2* (**G**) *Gapdh*, (**H**) *Rps10*, and (**I**) *Vegfa* expression relative to *Hprt*. n=5 biological replicates. Two-way ANOVA with Sidak’s multiple comparisons test. \**p*<0.05, \*\**p*<0.01, \*\*\**p*<0.001. Error bars represent standard deviation.

Gene transcription is not universally altered because there was no change in the expression of the housekeeping genes *Gapdh* and *Rps10* (**Fig. 3G-H**). Moreover, we did not detect any changes in *Vegfa* transcript levels, even though HEXIM1 is reported to modulate VEGF activity outside the nervous system (55, 56) (**Fig. 3I**). These findings show that the overexpression of HEXIM1 protein expression in neurons blunts IEG induction without effects on housekeeping genes or *Vegfa* and indicate that HEXIM1 can play a direct role in modulating IEG inducibility.

### The P-TEFb is required for IEG induction

HEXIM1 in other cell types helps suppress inducible gene expression through inhibiting P-TEFb activity (15, 57, 58), so we tested if small molecule inhibitors of P-TEFb influence immediate early gene activation in neurons. Inhibiting P-TEFb with 5,6-Dichloro-1-β-D-ribofuranosylbenzimidazole (DRB) leads to suppressed activation of *Fos* (**Fig. 4A**), *Arc* (**Fig. 4B**), *Egr1* (**Fig. 4C**), and *Nr4a2* (**Fig. 4D**) relative to cells treated with DMSO vehicle control. Similar results were obtained using a second structurally dissimilar and more potent P-TEFb inhibitor, Flavopiridol (FLAVO) (**Fig. 4E-H**) (59–61). These inhibitors similarly influenced activation of *Fos* and *Egr1* by KCl in dN2a cells (**Supp. Fig. 3A-D**). Furthermore, we tested the effects of a third highly selective inhibitor for CDK9 over other CDKs (62), JSH-009 (also known as Tambiciclib or GFH009). To identify an optimal concentration in our cells, we tested the effects of JSH-009 on the phosphorylation of a known CDK9 substrate, RNAP2-pS2. Other kinases phosphorylate RNAP2 at serine 5 (RNAP2-pS5). At 100nM, JSH-009 substantially reduced RNAP2-pS2 with little impact on RNAP2-pS5 in primary hippocampal neurons (**Fig. 4I**) and in dN2a cells (**Supp. Fig. 3E-G**). This concentration of JSH-009 attenuated KCl-dependent IEG activation in primary neurons (**Fig. 4J-M**) and in dN2a cells (**Supp. Fig. 3H-I**), like the effects of DRB and flavopiridol. Since basal IEG expression (in the absence of KCl) was affected by some of the CDK9 inhibitors (**Fig. 4J,L, Supp. Fig. 3A,D,H**), we tested for more global effects on gene expression. JSH-009 did not significantly change the expression of *Rps10 or Tcf4* (63) in primary neurons (**Fig. 4N-O**) or dN2a cells (**Supp. Fig. 3J-K**). However, *Vegfa expression* was significantly decreased by 100 nM JSH-009 in stimulated primary neurons (**Fig. 4P**) and in unstimulated dN2a cells (**Supp. Fig. 4L**). Overall, these data indicate that P-TEFb modulates stimulus-dependent IEG induction in neurons.

**Figure 4.**
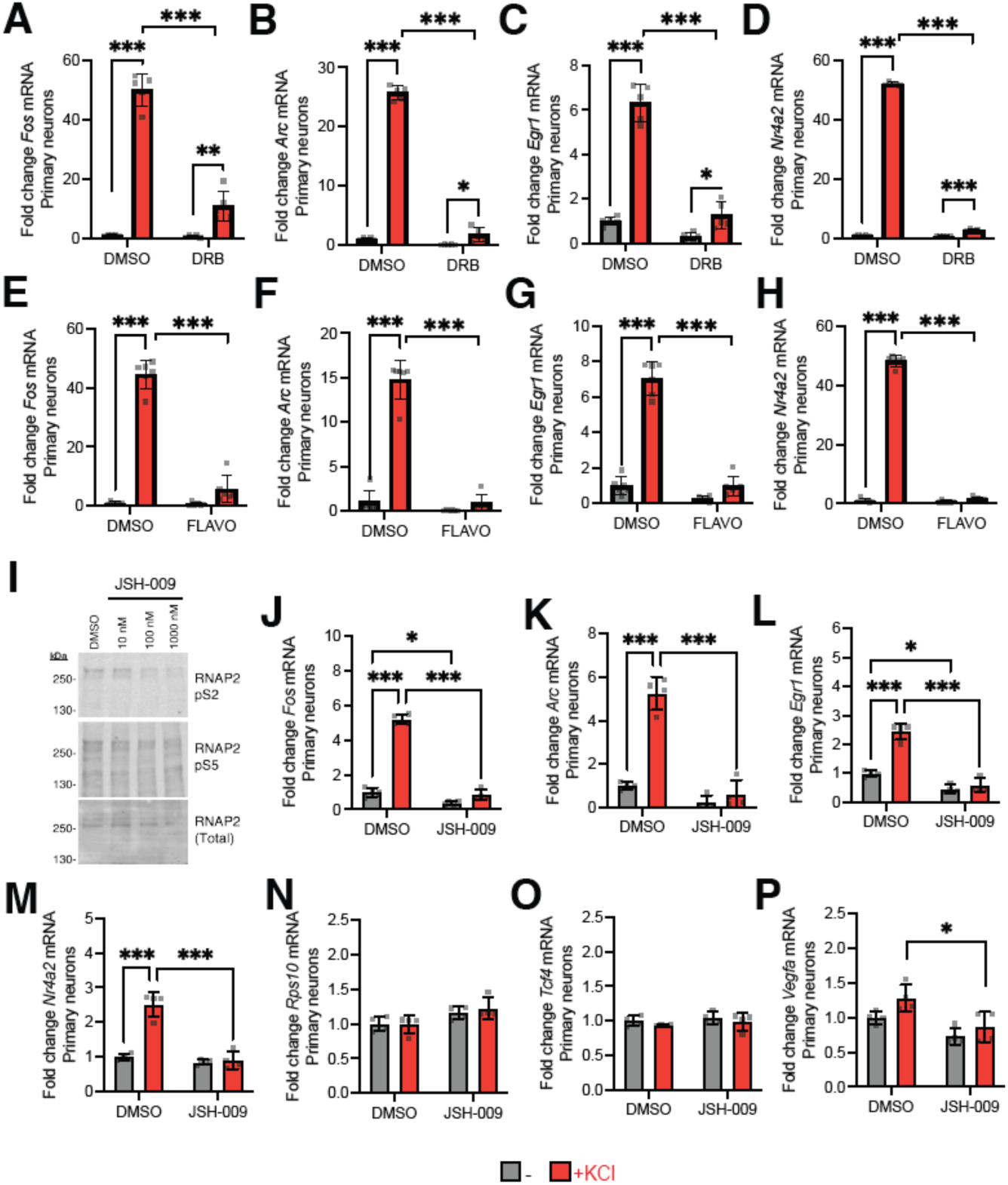
P-TEFb kinase CDK9 regulates activity-dependent IEG induction in hippocampal neurons. IEG mRNA levels (RT-PCR) were measured following P-TEFb inhibitor pretreatment then stimulation by 2 hr KCl in primary neurons. Average gene expression changes in (**A**) *Fos,* (**B**) *Arc*, (**C**) *Egr1*, (**D**) *Nr4a2* expression after DRB and KCl stimulation. Average gene expression changes in (**E**) *Fos,* (**F**) *Arc*, (**G**) *Egr1*, (**H**) *Nr4a2* after FLAVO and KCl stimulation. (**I**) Western blots of RNAP2-pS2 (top), RNAP2-pS5 (middle), and total RNAP2 (bottom) using whole cell primary hippocampal neuron lysates after 2.5 hr treatment with indicated doses of JSH-009. Average gene expression changes in (**J**) *Fos,* (**K**) *Arc*, (**L**) *Egr1*, (**M**) *Nr4a2*, (**N**) *Rps10*, (**O**) *Tcf4*, and (**P**) *Vegfa* after JSH-009 and KCl stimulation. All fold changes are calculated relative to *Hprt*. **A**-**D** n=5 biological replicates, **E**-**H** n=6 biological replicates, and **J**-**P** n = 4 biological replicates. Two-way ANOVA with Sidak’s multiple comparisons test. \**p*<0.05, \*\**p*<0.01, \*\*\**p*<0.001. Error bars represent standard deviation.

### HEXIM1/P-TEFb interaction influences IEG expression, and depends on calcium

We next tested whether HEXIM1/P-TEFb form a complex in neurons. We found that CDK9, the kinase subunit of P-TEFb, coimmunoprecipitates with HEXIM1 from primary cortical neuron protein lysates and vice versa (**Fig. 5A**). Both the 42 kDa and the 55 kDa isoforms (64) of CDK9 associate with HEXIM1. We further blotted for CCNT1 as an additional component of P-TEFb, and found it also associated with HEXIM1 and CDK9 in neurons. IgG negative control did not pull down these proteins. These lysates were generated using cortical rather than hippocampal primary cultures to ensure we had enough material to conduct this experiment as they are more abundant. Therefore, we also wanted to test for this interaction in the hippocampus. We generated protein lysates using whole adult mouse hippocampi and repeated the coimmunoprecipitations. Similarly, we observed that HEXIM1 associated with both isoforms of CDK9, and CDK9 associated with HEXIM1 (**Supp. Fig. 4A**). These data confirm that HEXIM1 interacts with P-TEFb in neurons and in the hippocampus.

**Figure 5.**
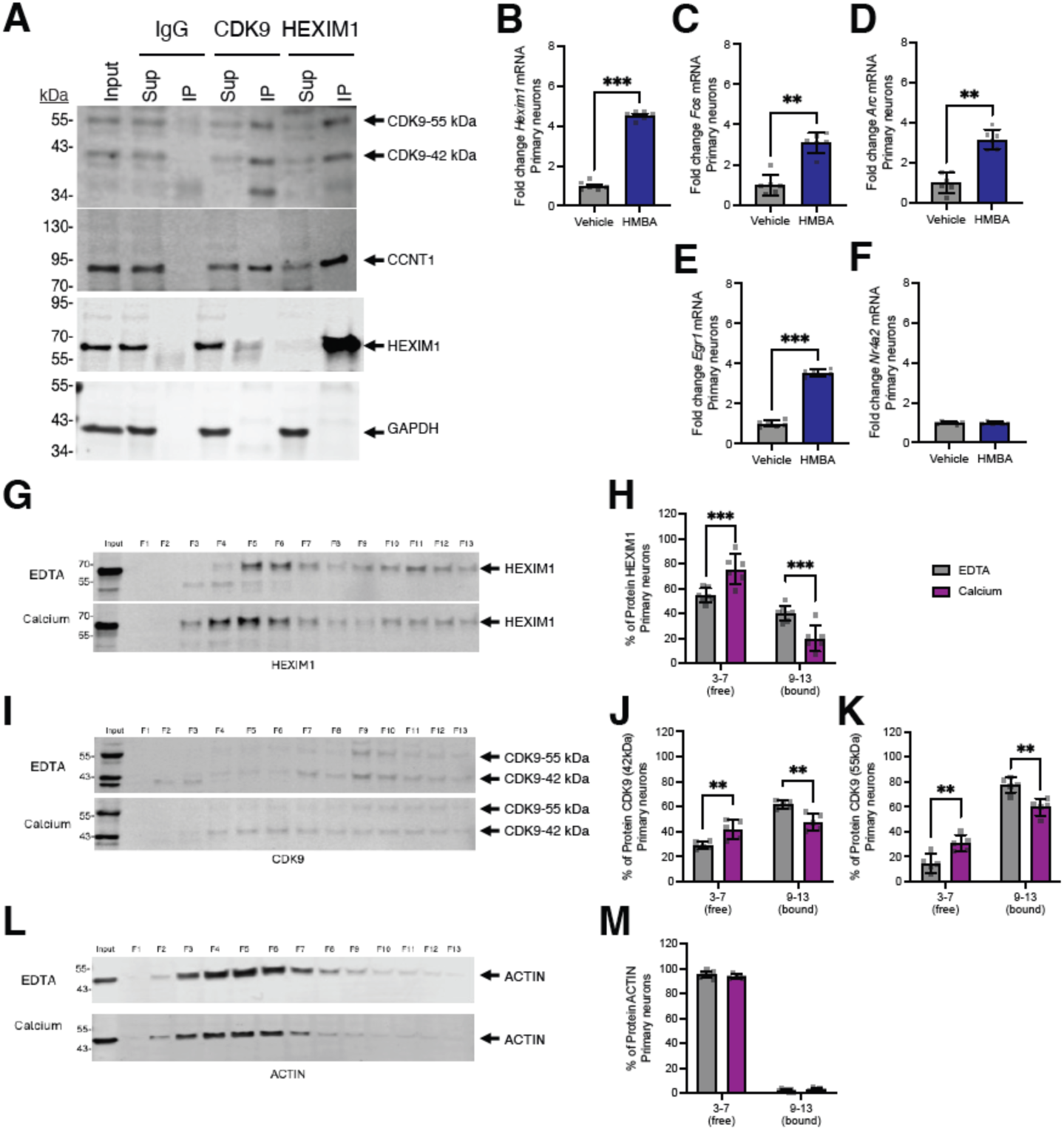
HEXIM1/P-TEFb binding and dissociation in primary neurons. (**A**) Co-immunoprecipitations from primary cortical neuron lysates with HEXIM1, CDK9, and IgG antibodies. Sup = supernatant and IP = co-immunoprecipitation. (**B-F**) Changes in gene expression following HMBA treatment in primary hippocampal neurons (RT-PCR). We depict (**B**) *Hexim1*, (**C**) *Fos*, (**D**) *Arc*, (**E**) *Egr1*, (**F**) *Nr4a2* mRNAs relative to *Hprt* mRNA. n=6 biological replicates. Paired two-tailed t test. (**G-M**) Representative blot of glycerol fractionation and quantitation of the percent of signal in low molecular weight fractions 3-7 and high molecular weight fractions 9-13 for each protein for (**G-H**) HEXIM1, (**J-K**) CDK9, and (**L-M**) ACTIN. For blots, input is shown in the far-left lane, then we loaded glycerol gradient fractions taken from the top of the column to the bottom. Thus, the lower molecular weight fractions are on the left and higher molecular weight fractions on the right. Two-way ANOVA with Sidak’s multiple comparisons test. n=6-7 biological replicates for HEXIM1 and ACTIN and n=5 biological replicates for CDK9 glycerol gradient quantitation. \*\**p*<0.01, \*\*\**p*<0.001. Error bars represent standard deviation.

Hexamethylene bisacetamide (HMBA) transiently dissociates HEXIM1 from P-TEFb (65, 66), freeing P-TEFb to induce transcription elongation unrestricted by the inhibitory complex (**Fig. 5A**). Therefore, we investigated whether the disruption of P-TEFb sequestration by HEXIM1 using HMBA affects IEG expression. As reported in other cell types, short-term HMBA treatment increased *Hexim1* mRNA expression in neurons (**Fig. 5B**) and dN2a cells (**Supp. Fig. 4B**) with no effect on HEXIM1 protein levels yet (**Supp. Fig. 4C**). HMBA treatment partially releases HEXIM1 from the high molecular weight P-TEFb complex in dN2a cells as separated using glycerol gradients (**Supp. Fig. 4D-E**), with a corresponding increase in the amount of HEXIM1 in low molecular weight fractions that contain actin. Interestingly, unlike previously published findings, CDK9 is still detected strongly in the high molecular weight complex (**Supp. Fig. 4D,F**). ACTIN distribution in the gradients was unaffected by HMBA (**Supp. Fig. 4D,G**). We then tested the effect of HMBA on IEG expression under basal conditions. Expression levels of *Fos* (**Fig. 5C; Supp. Fig. 4H**), *Arc* (**Fig. 5D**), and *Egr1* (**Fig. 5E; Supp. Fig. 4I**), were significantly increased by HMBA. However, the expression level of *Nr4a2* in primary neurons (**Fig. 5F**) was unchanged. These data suggest that when the HEXIM1 interaction with the high molecular weight complex is diminished, some IEGs are activated.

Since intracellular calcium is increased following depolarization, we next investigated the effect of adding calcium (1 mM) to cellular lysates from primary cortical neurons and assaying for changes in the size of P-TEFb complexes. Glycerol gradients with and without calcium were run. The no calcium control lysates also contained EDTA to chelate free calcium. Calcium addition partially disrupts the P-TEFb complex, as shown by reciprocal changes in the amounts of both HEXIM1 and both isoforms of CDK9 in low and high molecular weight fractions (**Fig. 5G-K**), without changes to the distribution of ACTIN (**Fig. 5L-M**). These findings suggest that increased of intracellular calcium might directly induce dissociation of the P-TEFb complex.

### Role of HEXIM1/P-TEFb interaction at gene promoters promotes Fos transcription

The function and regulation of P-TEFb at IEG promoters in neurons is not well defined. Moreover, while HMBA is reported to dissociate the inhibitory P-TEFb complex, the specificity of HMBA is unclear (67, 68), so we wanted to investigate the impact of the HEXIM1/P-TEFb complex interaction on KCl induced gene expression with a more direct approach. First, we examined CDK9 binding at the *Fos* promoter in primary neurons in a published dataset that also utilized KCl stimulation (69). This dataset indicates that CDK9 is bound to the *Fos* promoter regardless of whether neurons are at baseline or depolarized (**Fig. 6A**). Moreover, an experiment in a human melanoma cell line both suggests CDK9 and HEXIM1 co-occupy the *Fos* promoter (**Supp Fig. 5A**). We therefore wondered if HEXIM1 is also present at the *Fos* promoter in our system. We conducted chromatin immunoprecipitation (ChIP) with HEXIM1 antibody, then probed the CDK9 binding site using quantitative PCR (primer target position indicated on the bottom of **Fig. 6A**). We observed about 4-fold enrichment of HEXIM1 at the *Fos* promoter in unstimulated N2a cells relative to unenriched input samples (**Fig. 6B**). Together, these experiments indicate that the inactive P-TEFb complex is bound to the *Fos* promoter under basal conditions alongside HEXIM1, possibly poising the gene for P-TEFb-dependent activation.

**Figure 6.**
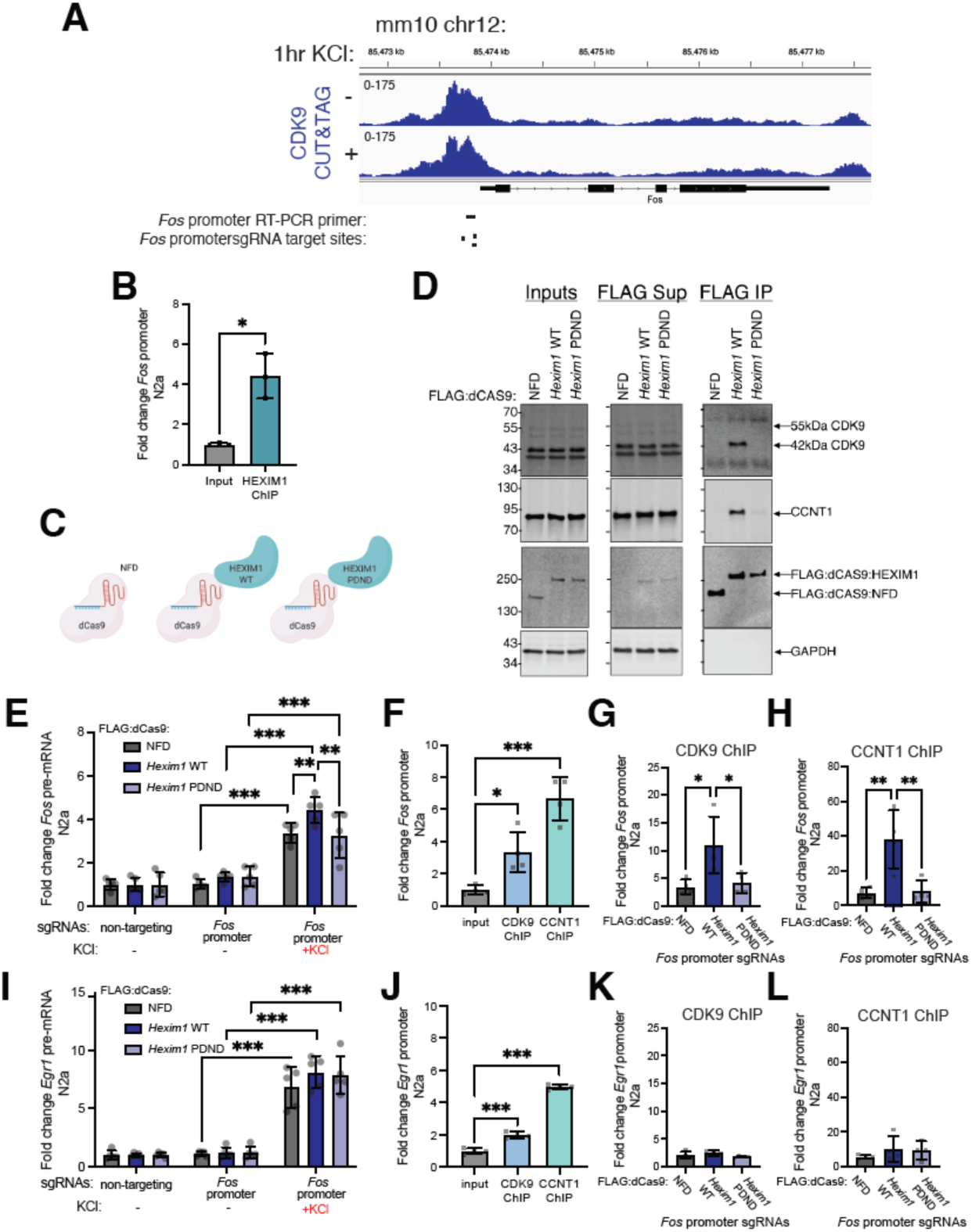
Targeting HEXIM1 to *Fos* promoter activates *Fos* transcription via its interactions with P-TEFb. (**A**) CDK9 signal from CUT&TAG experiment in primary mouse neurons without KCl stimulation or with KCl stimulation at *Fos* promoter. Locations of ChIP primer target location and sgRNA targets are indicated underneath gene tracks. (**B**) Analysis of HEXIM1 enrichment by ChIP relative to input chromatin at the *Fos* promoter in unstimulated N2a cells. Signal is normalized to intronic region of *Hdac2* gene. Paired two-tailed t test (n=3). (**C**) *Hexim1* was cloned such that the expressed protein is tethered to a nuclease dead Cas9. Controls include a dCas9 with no functional domain attached (the NFD construct) and HEXIM1 with a mutation of its PYNT domain (that interacts with P-TEFb) to PDND. (**D**) Coimmunoprecipitations were conducted to test P-TEFb interactions of the three dCas9 proteins in dN2a. Western blots of inputs, supernatant (Sup), or FLAG immunoprecipitation (IP) are depicted for CDK9, CCNT1, FLAG tag, and GAPDH top to bottom. All cells were co-transfected with *Fos*-promoter targeting guide RNAs. (**E**) Following transfection, cells were stimulated with KCl (50mM for 90 minutes). The respective changes in *Fos* pre-mRNA in N2a were detected by RT-PCR. Fold changes are relative to *Hprt*. n=5 biological replicates. Two-way ANOVA with Tukey multiple comparisons test. (**F**) qPCR of a *Fos* promoter region in CDK9, and CCNT1 ChIPs relative to purified input chromatin in dCas9-NFD control samples with *Fos* promoter sgRNAs. Significance of enrichment for each ChIP relative to the input at the *Fos* promoter normalized to intronic region of *Hdac2* gene was determined with one-way ANOVA followed by Dunnett’s multiple comparisons test. (n=4). (**G**-**H**) ChIP-qPCR analysis of (**G**) CDK9 or (**H**) CCNT1 at the *Fos* promoter normalized to intronic region of *Hdac2* gene. One-way ANOVA with Tukey’s multiple comparisons test (n=4). (**I**-**L**) Same analysis as described for *Fos* transcription and promoter binding in **E**-**H**, but this time examining (**I**) *Egr1* pre-mRNA expression changes and (**J**-**L**) *Egr1* promoter binding when the dCas9 constructs are targeted to the *Fos* promoter. \**p*<0.05, \*\**p*<0.01, \*\*\**p*<0.001. Error bars represent standard deviation.

We next probed the function of HEXIM1 at the *Fos* promoter with a genomic targeting strategy. In combination with guide RNAs, nuclease dead CRISPR associated protein 9 (dCas9) can target an epigenetic regulator to specific sites in the genome without cutting the DNA. Therefore, we used a set of three dCas9 expression constructs: a control with just the dCas9 with no functional domain (dCas9-NFD), HEXIM1 wild-type (WT; dCas9-HEXIM1 WT), and a HEXIM1 mutant that disrupts the PYNT motif responsible for HEXIM1’s binding to P-TEFb (dCas9-HEXIM1 PDND) (**Fig. 6C**). We designed guide RNAs to hybridize at the *Fos* promoter near where CDK9 is bound (position indicated on bottom of **Fig. 6A**). To first test that the guides we designed can effectively target an epigenetic regulator to the *Fos* promoter, we used dCas9 tethered to the catalytic core of the transcription activator p300. Relative to cells transfected with non-targeting guides that contain sequences not matching any sites within the mouse genome, directing binding of the p300 core to the *Fos* promoter significantly activated *Fos* mRNA expression (**Supp Fig. 5B**). The PYNT domain is responsible for the binding of HEXIM1 with P-TEFb. We tested if mutation of this domain alters binding of HEXIM1 with P-TEFb as previously reported (70) by conducting CoIPs using the FLAG tag on the dCas9 constructs. While CDK9 and CCNT1 efficiently co-precipitated with dCas9-HEXIM1 WT, the PDND mutation largely disrupted this interaction (**Fig. 6D**). We identified transfection conditions that produced limited dCas9-HEXIM1 expression relative to endogenous HEXIM1 protein (**Supp Fig. 5C**) to avoid effects of overexpression on KCl-stimulated transcription. We confirmed that the change in total HEXIM1 levels was not sufficient to impact KCl activation if dCas9 was not targeted to a specific genomic locus (**Supp Fig. 5D-E**). We were surprised to find that basal expression of the *Fos* pre-mRNA was unaffected by the targeting of HEXIM1 WT or PDND to the *Fos* promoter. However, targeting HEXIM1 WT to the *Fos* promoter significantly enhanced *Fos* transcription activation in response to KCl-induced depolarization, and this effect was abolished by the PDND mutant (**Fig. 6D**). We posited that perhaps HEXIM1 WT was helping recruit P-TEFb at baseline, but keeps it inactive until cells are depolarized. Therefore, we tested for changes in P-TEFb recruitment in unstimulated cells using ChIP when the dCas9 constructs are targeted to the *Fos* promoter. We detected significant enrichment of CDK9 and CCNT1 at this site relative to Input (**Fig. 6F**) in the dCas9-NFD control cells, suggesting P-TEFb is enriched at the *Fos* promoter in N2a cells. Moreover, when HEXIM1 WT is recruited to the promoter by the dCas9 system, CDK9 (**Fig. 6G**) and CCNT1 (**Fig. 6H**) are significantly enriched at the same site relative to dCas9-NFD and dCas9-HEXIM1-PDND controls. We also tested the specificity of these transfections by examining *Egr1* pre-mRNA expression when the *Fos* promoter was being targeted. While KCl induced *Egr1* pre-mRNA expression, we observed no differences in *Egr1* expression between dCas9-NFD, dCas9-HEXIM1 WT, or dCas9-HEXIM1 PDND groups in unstimulated or stimulated conditions (**Fig. 6I**). Like at the *Fos* promoter, CDK9 and CCNT1 are also significantly enriched at the *Egr1* promoter in the NFD control ChIP samples (**Fig. 6J**). However, the recruitment of HEXIM1 to the *Fos* promoter with dCas9 had no effect on CDK9 (**Fig. 6K**) or CCNT1 (**Fig. 6L**) enrichment at the *Egr1* promoter. Together, this set of experiments suggests that in the absence of a stimulus, HEXIM1 helps recruit silenced P-TEFb to the *Fos* promoter to poise it for activation. Following stimulation, however, HEXIM1 can release the inhibition of P-TEFb to activate IEG transcription.

### HEXIM1 protein is diminished, and induction of some IEGs is suppressed, for several hours following neuronal depolarization

We predict that re-establishment of the basal state after a depolarization requires the re-sequestration of P-TEFb by HEXIM1 in the inhibited complex. Therefore, we first investigated the impact of depolarization on HEXIM1 protein levels in primary neurons. Immediately after a 2 hr. KCl stimulation, HEXIM1 and CDK9 levels are not significantly changed (**Fig. 7A-D**). However, we reasoned that the increased *Hexim1* mRNA expression might have effects on HEXIM1 protein levels at later time points after depolarization. Therefore, we stimulated cells with KCl for 2 hrs., then removed the stimulation for 4-24 hrs, examining HEXIM1 expression at these times (**Fig. 7E**). We found that HEXIM1 protein levels are surprisingly decreased 4 and 6 hr after depolarization and recover back to baseline levels after 24 hrs (**Fig. 7F-G**).

**Figure 7.**
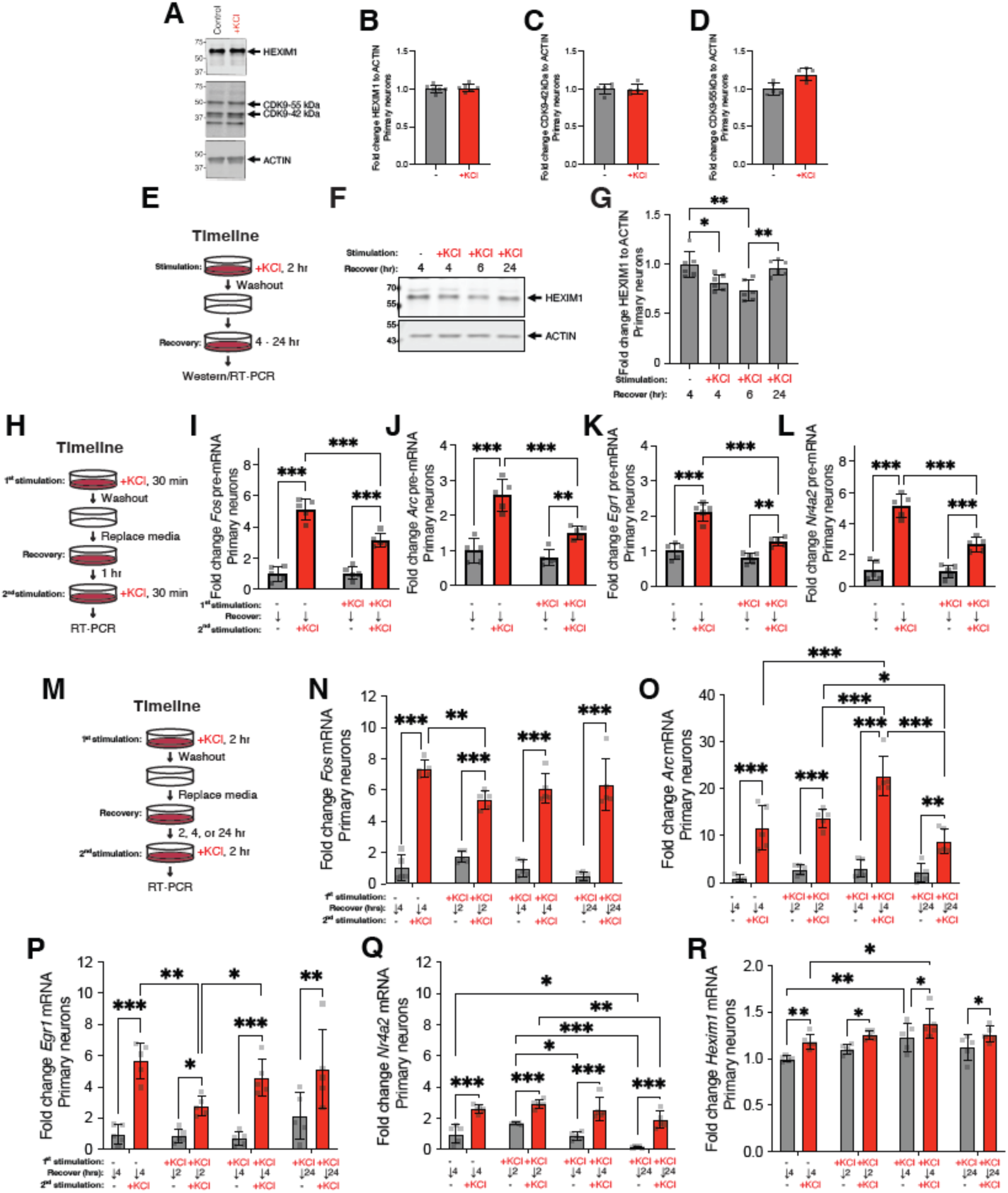
HEXIM1 protein in primary hippocampal neurons following KCl stimulation, and transcriptional responses of IEGs and *Hexim1* after a prior stimulation. (**A**) Representative Western blots of HEXIM1 and CDK9 relative to ACTIN after KCl stimulation. (**B-D**) Quantitation of proteins expression relative to ACTIN for (**B**) HEXIM1, (**C**) CDK9 42kDa, and (**D**) CDK9 55kDa after KCl across n=6 biological replicates. Paired two-tailed t test did not detect statistical differences between conditions for **B**-**D**. (**E**) Timeline of KCl stimulation and recovery in **F** and **G**. (**F**) Representative Western blot of HEXIM1 relative to ACTIN after KCl stimulation and washout. (**G**) Quantitation of HEXIM1 after KCl across n=6 biological replicates. One-way ANOVA with Tukey multiple comparisons tests. (**H**) Timeline of briefer KCl stimulations and restimulations applied to samples depicted in **I-L**. (**I-L**) pre-mRNA expression (RT-PCR) of IEGs after KCl stimulations. The genes (**I**) *Fos*, (**J**) *Arc*, (**K**) *Egr1*, and (**L**) *Nr4a2* pre-mRNA expression is depicted. Two-way ANOVA with Sidak’s multiple comparisons test. (**M**) Timeline of longer KCl stimulations, longer recovery, and restimulations in **N-R**. (**N-R**) mRNA expression (RT-PCR) after treatments following this timeline. The genes (**N**) *Fos*, (**O**) *Arc*, (**P**) *Egr1*, (**Q**) *Nr4a2*, and (**R**) *Hexim1* mRNA expression are depicted. All experiments conducted in primary hippocampal neurons and fold changes are calculated relative to *Hprt*. Two-way ANOVA with Tukey multiple comparisons test. n=5 biological replicates. \**p*<0.05, \*\**p*<0.01, \*\*\**p*<0.001. Error bars represent standard deviation.

We wondered if the decrease in HEXIM1 correlates with differences in transcriptional responses to a second KCl. Therefore, to directly test whether depolarization results in a short-term suppression of transcriptional responses to a second stimulation, we stimulated cells with KCl for 30 min, washed out the KCl and allowed them recover for 1 hr, and then re-stimulated them with KCl for 30 min (**Fig. 7H**). On these short time scales, IEG expression was measured using primers directed to pre-mRNA transcripts. Basal transcription of the four IEGs recovered to near-normal levels 1 hour after the KCl washout. However, transcriptional responses to the second depolarization at the *Fos*, *Arc*, *Egr1*, and *Nr4a2* genes were significantly suppressed (**Fig. 7I-L**). Dampening of several IEGs was previously observed with a 1 hr window between stimulations (10). However, because we saw HEXIM1 protein levels were reduced several hours after an initial stimulation, we asked if recovery of transcriptional responses may take longer than 1 hr., even though mitogen-activated protein kinase signaling pathways, as measured by phosphorylated extracellular signal–regulated kinase, were reported to be recovered after 2 hrs. (10). Therefore, we examined the temporal dynamics of transcriptional repression and recovery over longer time scales using mRNA primers (**Fig. 7M**). We found that *Fos* mRNA expression to a second depolarization was still significantly suppressed after a 2 hr recovery period compared to the depolarization of naïve cells. However, the transcriptional response of the *Fos* gene recovered to near-normal levels after a 4 or 24 hr recovery period (**Fig. 7N**). Transcription of *Egr1* was similarly suppressed after 2 hr, and recovered to near-normal levels after 4 or 24 hrs (**Fig. 7P**). Transcriptional responses of *Arc* and *Nr4a2* to the second depolarization were not significantly suppressed after a 2 hr recovery (**Fig. 7O, 7Q**). Moreover, after a 4 hr recovery, we detected an enhanced transcriptional response to the second depolarization at the *Arc* gene, which normalized after a 24 hr recovery (**Fig. 7O**). In contrast, we detected significant changes mostly in baseline transcription of *Nr4a2* using the same stimulation-recovery-re-stimulation paradigm (**Fig. 7Q**). Parallel studies in dN2a cells revealed a similar transient suppression and recovery of *Fos* and *Egr1* expression (**Supp. Fig. 6A-B**). In parallel, we observed reciprocal changes in the levels of *Hexim1* mRNA after KCl treatment, and see further increases in its transcription upon repeated stimulations (**Fig. 7R**; **Supp. Fig. 6C**), suggesting that cells might synthesize more *Hexim1* mRNA to restore HEXIM1 protein levels by 24 hours to re-establish the normal basal levels of poised RNAP2 to be prepared for another round of IEG induction.

### Blocking P-TEFb prevents depolarization-induced suppression of IEG induction

The transient suppression and recovery of *Fos* and *Egr1* transcription in response to repeated depolarization is consistent with the potential involvement of HEXIM1 and P-TEFb in regulating the pause/release of RNAP2 at these genes. To test if P-TEFb activity modulates responses to repeated depolarization, we tested whether inhibiting P-TEFb using DRB during the first KCl stimulation prevents the transient suppression of transcription in response to a second stimulation. DRB was chosen as the P-TEFb inhibitor for this experiment rather than FLAVO or JSH-009 because of its known ability to be efficiently washed away (71). The cells were thoroughly washed after the first stimulation to remove both DRB and KCl, and cells were allowed to recover for 2 hrs before being re-stimulated with KCl in the absence of DRB (**Fig. 8A**). Transcription of *Fos* (**Fig. 8B**) and *Egr1* (**Fig. 8C**) mRNA following a second depolarization was suppressed, as we saw previously. However, the presence of DRB during the first stimulation significantly alleviated the suppression of transcriptional responses in the second stimulation, with *Fos* almost fully recovering to normal induction levels. The interpretation of these data may be confounded if effects of the transient DRB treatment persisted after the recovery period, despite the prior washout. Therefore, in additional control studies, cells were treated with DMSO or DRB alone, washed and allowed to recover for 2 hr and then depolarized. The initial DRB treatment had no effect on the basal expression of either *Fos* or *Egr1*, or on the depolarization-induced expression of *Fos* (**Supp. Fig. 7A**). However, the depolarization-induced expression of *Egr1* was partially suppressed by the initial DRB treatment, despite the washout (**Supp. Fig. 7B**). Therefore, the incomplete rescue of *Egr1* expression by DRB pre-treatment in **Fig. 8C** could be due to residual effects of DRB on *Egr1* transcription after the washout. Overall, these results indicate that P-TEFb releases paused RNAP2 during the first stimulation, leading to transient suppression of responses to a second stimulation because RNAP2 is not already paused on the IEG to facilitate a naïve level of response to a second stimulation.

**Figure 8.**
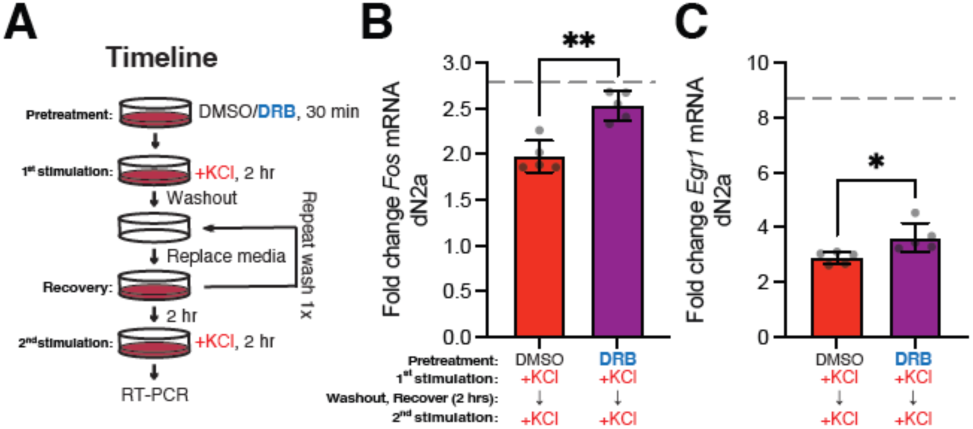
Blocking P-TEFb preserves KCl induction of *Fos* and partially recovers *Egr1* inducibility during a second stimulation. **(A)** Timeline of drug treatment and KCl stimulations in dN2a cells. (**B-C**) RT-PCR of gene expression changes following two successive KCl stimulations with or without DRB pretreatment prior to and during the first stimulation. The expression of (**B**) *Fos* and (**C**) *Egr1* mRNA is depicted. Effect of KCl alone (done at same time as the second stimulation) with no prior drug or KCl treatments is shown as a dashed line. n=5 biological replicates. Paired two-tailed t test. * = p < 0.05, \*\**p*<0.01. Error bars represent standard deviation.

## Discussion

The regulation of P-TEFb has been linked to multiple brain diseases and disorders (72–74), but little is known about the role of P-TEFb regulators in AD. *HEXIM1* was previously suggested to be dysregulated in AD (75), and we show here that *HEXIM1* expression is correlated with worse neuropathology and cognition. Tau pathology and longitudinal cognition in neurons have the strongest associations in individual cell subtypes detected by snRNA-seq. We also highlight that increased *HEXIM1* expression is found in excitatory neurons in human AD across several brain regions in a separate snRNA-seq cohort (52). Inhibitors that block RNAP2 pause release have demonstrated therapeutic effects in AD animal models (30–34, 40–44). The increased *HEXIM1* mRNA levels in human AD neurons might suggest transcription pause release is suppressed, a seemingly paradoxical finding. However, there are several potential explanations. First, blocking pause release via HEXIM1 upregulation could be a protective response of the brain to AD pathology, so furthering this block with pause release inhibitors would be beneficial to memory function. Second, we do not yet know if the HEXIM1 protein is increased or decreased in AD. If protein levels are decreased, *HEXIM1* mRNA could be increased in a compensatory attempt to make more HEXIM1 protein, as we observed following our KCl stimulations. This decrease in HEXIM1 protein could allow for overly permissive elongation that is corrected by inhibiting transcriptional pause release. Despite not yet fully understanding the role of HEXIM1 in AD, our findings indicate that HEXIM1 dysregulation in either direction could affect IEG inducibility, and this could create a possible mechanistic link explaining cognitive impairments in AD, as IEGs are crucial to memory formation (76–80). Together, these observations indicate that it is important to develop a better understanding of how HEXIM1 is involved in regulating neuronal gene transcription.

We observed that *Hexim1* mRNA is increased in the hippocampus during memory formation. Unbiased interrogations of mRNA expression changes after fear stimulus or novel environment exposure have identified upregulation of Hexim1 mRNA expression in mice and rats as well (81–83), suggesting a variety of stimuli induce this gene’s expression in multiple species. Based on the results of our time course examining HEXIM1 protein levels after KCl stimulation, we predict that this upregulation in *Hexim1* mRNA could help neurons replenish HEXIM1 protein to baseline levels, though additional testing of this hypothesis is required.

Although HEXIM1 has been implicated as a key regulator of P-TEFb, the role of P-TEFb regulation in neurons is poorly understood. In this work we found that inhibiting P-TEFb with DRB, FLAVO, or JSH-009 blocked stimulus-dependent activation of the four IEGs we tested, consistent with prior studies of *Arc* and *Fos* regulation (11, 79) and demonstrating that P-TEFb likely plays a central role in stimulus-dependent activation of neuronal IEGs. P-TEFb might also influence the expression of *Vegfa*, which is important for a variety of neuronal functions and protective against neurodegeneration (84).

Although JSH-009 has significantly improved specificity over FLAVO and DRB, a characterization of its off-target effects suggest it still has some activity towards dual specificity tyrosine-phosphorylation-regulated kinase 1 A and B (DYRK1A and DYRK1B) (62), and these kinases are also inhibited by FLAVO (85). Therefore, more specific manipulations are required to disambiguate the effects of CDK9 from DYRK1A and DYRK1B on IEG activation in neurons.

The interaction between HEXIM1 and P-TEFb seems particularly important for IEG modulation. Evidence for this includes our findings that overexpression of HEXIM1 alters IEG induction, HMBA treatment that frees HEXIM1 activates IEGs, calcium dissociates P-TEFb from HEXIM1, and a mutation that impairs P-TEFb binding dampens HEXIM1-facilitated *Fos* activation. In its free form in the cell, HEXIM1 overexpression blocks IEG activation following KCl treatment, suggesting P-TEFb sequestration is too strong in neurons when HEXIM1. Moreover, HMBA release of HEXIM1 from the repressive complex increases IEG expression. We do not know precicely why CDK9 remains in the higher molecular weight fractions after HMBA treatment of cells, but there are a few possibilities. Perhaps only HEXIM1 leaves the complex in neuronal cells. If this is the case, we think this might be sufficient for P-TEFb activation, as a peptide sequence within HEXIM1 directly modulates access to the CDK9 catalytic cleft (86). It is also possible that it becomes associated with another high molecular weight complex, like the superelongation complex (65) since glycerol gradients can only distinguish associations with complexes of different sizes. However, our gradients showed that calcium releases both HEXIM1 and CDK9 into lower molecular weight complexes. It is still unclear how calcium is affecting the complex. It could bind a component of the complex directly, alter calcium-dependent phosphorylation, or could be mediated by another calcium-binding protein, such as calmodulin. Finally, our dCas9 tethering experiments combined with published CDK9 CUT&TAG experiments, suggest that focal binding of HEXIM1 at IEG promoters may help recruit and sequester P-TEFb in an inactive form at gene promoters, helping to poise genes for activation. During depolarization, increases in intracellular calcium could allow for disinhibition of P-TEFb at gene promoters to activate IEG expression.

Since HEXIM1 protein levels decrease during recovery from an initial depolarization for an even longer time than the recovery of some calcium signaling pathways (10) we hypothesized that the normal sequestration of P-TEFb is impaired following depolarization, consistent with the reduced induction of some IEGs as previously observed (10, 87). We further observe that this suppression persists for several hours for some genes. The recovery of the transcriptional response of the *Fos* and *Egr1* genes takes several hours and seems to specifically involve P-TEFb. Interestingly, learning is dependent on temporal spacing of training. For example, massed training paradigms with less than 1 hr intervals between training session show impaired long-term recall compared to intervals spread out across days (88, 89). However, intervals of several hours on the same day show long-term memory enhancement (90, 91). Further work is required to determine if the transcriptional impairment observed with closely-spaced stimuli underlie the impaired plasticity and memory at similar timescales. By contrast, while transcription of *Arc* and *Nr4a2* is initially suppressed following depolarization, transcription appears to be fully recovered to control levels within 2 hours. We hypothesize that very rapid recovery of *Arc* expression leading to an enhanced transcriptional response to a second depolarization may reflect its unique role among these IEGs in encoding a cytoskeletal protein directly implicated in synaptic plasticity (92). *Nr4a2* also exhibits altered sensitivity to HMBA, suggesting it might be regulated by different regulatory mechanism(s) than other IEGs. Taken together, our data indicate that different IEGs exhibit different sensitivities to HEXIM1/P-TEFb regulation at baseline and indicate that multiple mechanisms control the recovery of transcriptional responsiveness.

In conclusion, *HEXIM1* mRNA levels exhibit a strong correlation with poorer cognition in AD, particularly in excitatory neurons. This highlights the need to better understand its role in inducible gene expression in neurons. Importantly, we find that the HEXIM1 protein supports poising of IEGs at baseline so they may respond strongly and quickly to stimuli. This is a critical first step in understanding the molecular function of HEXIM1 in regulating the expression of memory-associated genes in neurons.

### Experimental procedures

#### ROS/MAP longitudinal study and autopsy

Studies were approved by an Institutional Review Board (IRB) of Rush University Medical Center, and comply with the Declaration of Helsinki principles. Participants free from known dementia enrolled and agreed to annual clinical evaluation and donation of their brain at the time of their death. All participants signed informed and repository consents and an Anatomic Gift Act for brain donation (93). Secondary analyses of this extant data were approved by the Vanderbilt University Medical Center IRB. Cognition was defined as a global cognition composite, an average of z-scores from 17 tests across 5 domains of cognition (episodic, semantic and working memory, perceptual orientation, and perceptual speed). The *z* scores of all the available tests were averaged to create a global cognition composite. Methodology for calculating cognitive scores were detailed previously (94). AD pathology was determined by average percent area occupied by Aβ_42_ or Tau (AT8 epitope) across eight brain regions at autopsy: hippocampus, angular gyrus, and entorhinal, mid frontal, inferior temporal, calcarine, anterior cingulate, and superior frontal cortices (95, 96). Values were transformed to approximate a normal distribution of pathology.

#### Bulk RNA-seq

Data was accessed via the AD Knowledge Portal (Accession number: syn23650893). Data sets include RNA-seq of bulk tissue from the caudate nucleus (CN), dorsolateral prefrontal cortex (DLPFC), and posterior cingulate cortex (PCC) brain regions. Regions were selected in the parent study based on tissue availability and biological relevance to several disease conditions. Reads were aligned to Ensembl human reference genome (GRCh38, v99) with STAR (V2.5.2b). featureCounts (v2.0.0) was used to determine read counts per gene and Picard metrics were calculated (v2.18.27). Exclusion criteria are detailed previously (97). Number of subjects and demographic information for this analysis is detailed in **Supplemental Table 1**. RNA-seq expression was analyzed for all genes encoding proteins identified as interactors of CDK9 using the STRING database (98) with the following parameters: network type = full STRING network; active interaction sources = textmining, experiments, and databases, minimum interaction score = 0.9, 1^st^ shell = non/query proteins only; excluded disconnected nodes.

#### snRNA-seq

snRNA-seq data from DLPFC was sourced from the ROS/MAP longitudinal study and autopsy (51). Data was accessed via the AD Knowledge Portal (Accession number: syn31512863). Exclusion criteria were previously detailed (94), resulting in analysis of 424 participants. Negative binomial lognormal mixed models implemented in the NEBULA R package (v1.2.0) were used for this analysis, using sex, age at death, postmortem interval and clinical group as covariates. Genes with expression in a minimum 10% of all cells were included in this model. Cells were removed if they counted more than 20,000 or less than 200 total RNA unique molecular identifiers (UMIs) or had more than 5% mitochondrially mapped reads. The gene count matrix input of the model was the UMI count data from RNA assay normalized and scaled by “sctransform” R package (https://github.com/satijalab/sctransform). NEBULA-HL method is used for the modeling. The p value calculated from the models was FDR adjusted using R function “p.adjust” using “BH” as the method.

Analysis of snRNA-seq across multiple brain regions was previously reported, and we downloaded the author’s analysis (52). Specimens from six brain areas (entorhinal cortex, hippocampus, anterior thalamus, angular gyrus, midtemporal cortex, and prefrontal cortex) were collected from 48 subjects and analyzed by the authors. Cognitive impairment was defined on a five-point scale and pathology was determined by NIA-Reagan scores on a four-point scale (52, 99). Differentially expressed genes (DEGs) were identified by two statistical analyses: model-based analysis of single-cell transcriptomics (MAST) and Nebula testing using a Poisson mixed-model (PMM). DEGs are defined when both analyses identify them as significant (P_adj_ < 0.05) and the fold change direction is consistent by both analyses. For our graphs, when P_adj_ = infinity, it was assigned the value of 400 for the purposes of plotting.

#### Fear conditioning RNA-seq

RNA-seq reads from wild-type mice were sourced from a previously published dataset (53). Reads were aligned to mm10 using hisat2-2.0.4 (100). featureCounts was used to identify reads associated with refseq annotated genes and calculate FPKM, and reads overlapping more than one meta-feature were included. The EdgeR (101) package was used to perform a Fisher’s exact test and p values underwent FDR correction. Average FPKM and FDR from right and left hemisphere datasets for each animal is reported. In all cases where FDR is reported as significant, FDR from each hemisphere was <0.001.

#### Animals

C57BL/6J mice were purchased from Envigo or the Jackson Laboratory. Mice were group housed, kept under 12:12 light/dark cycles, with food and water available *ad libitum*. All procedures were approved by the Vanderbilt or Loyola Institutional Animal Care and Use Committees and conducted in full compliance with the Association for Assessment and Accreditation of Laboratory Animal Care (AAALAC).

#### Cell Culture

Primary neuron cultures were made from neonatal (P0) mice. Dissected hippocampi or cortices were dissociated with papain supplemented with cysteine in Hank’s balanced salt solution, and triturated to dissociate neurons in neurobasal complete media (neurobasal with 1× B27 supplement, 1 mM sodium pyruvate, 1 mM HEPES, 100 U/mL penicillin, 100 μg/mL streptomycin, and 0.5 mM L-glutamine) plus 10% fetal bovine serum (FBS). Cells were passed through a 100-μm filter (Falcon) and cells were applied to a poly-d-lysine (PDL) coated plate in neurobasal complete media without FBS. On day *in vitro* 1 (DIV 1), media was again changed to neurobasal complete without FBS, and ½ the media was changed every 2-3 days thereafter. All primary neuron experiments were performed using neurons at DIV 12–15. Biological replicates were made from different preparations of primary neurons.

Neuro-2a (N2a) cells (CCL-131) were obtained from ATCC and maintained according to their recommended conditions. To differentiate N2a, cells were attached overnight in growth media, then media was changed to differentiation media (DMEM with L-glutamine without glucose, 10 mM galactose, 100 U/mL penicillin, 100 μg/mL streptomycin, and 1× N2 supplement). After 2 days, media was changed to neurobasal complete media. For drug/KCl studies, cells were incubated in neurobasal overnight prior to the application of drug or KCl. For the restimulation time course study, KCl was applied at least 1 hr after switching to neurobasal complete for the 24 hr washout time point, then the subsequent time points occurred on the next day. Biological replicates of N2a cells were prepared from different passages of the cell line.

#### KCl treatments

KCl was prepared in water and sterile filtered. Stimulations occurred for 30 min., 90 min., or 2 hrs, as indicated. Primary neurons were stimulated with 25mM and dN2a were stimulated with 50mM KCl. KCl was added directly to neurobasal media, and thus the indicated mM is in addition to the 5.33mM KCl present in the normal composition of neurobasal media.

#### dCas9/sgRNA transfection

For RNA and Western blot analysis, 30,000 N2a cells in 24-well plates were plated in growth media. The next day, cells were transfected with GenJet for Neuro-2A cells (SignaGen) according to manufacturer recommendations, except the transfection mix contained 0.5μL of GenJet, 125ng dCas9 construct, and 62.5ng of pooled sgRNA expression vectors total in 100μL of serum-free media. For CoIP, 150,000 N2a cells were plated in a 6-well plate, and 625ng of dCas9 and 312.5 pooled sgRNA were transfected with 2.5uL GenJet in 500uL of serum-free media. For ChIP, 100mm plates with 900,000 cells were transfected using 3.75μg dCas9 vector, 1.875μg sgRNA vector, and 15μL GenJet or LipoD293 (SignaGen) in 3mL of serum-free media. Two plates per condition were pooled to make a batch of chromatin. Cells were kept in growth media 1-day following transfection, then media was changed to neurobasal the afternoon before stimulating with KCl and harvesting protein or RNA.

#### RT-PCR

Total RNA was extracted from samples using RNeasy Plus Mini kit (QIAGEN) and eluted in 30–50 μl of RNase free water. Sample concentration was analyzed using a NanoDrop One Microvolume UV-Vis Spectrophotometer (Thermo Scientific) and cDNA synthesis was conducted in 10-μl reaction using SuperScript VILO master mix (Invitrogen) according to manufacturer instructions. All cDNA reactions were diluted 1:10 with RNase-free water. RT-PCR was performed on a BioRad CFX96 or Opus CFX384 RT-PCR detection system in 10-15 μl reactions containing SsoAdvanced Universal SYBR Green Supermix and 250 μM primer and 3-4.5 μL of diluted cDNA. Relative fold quantification of gene expression between samples was calculated using the comparative C_t_ method (102) and normalized to *Hypoxanthine phosphoribosyltransferase* (*Hprt*). Primer sequences are listed below.

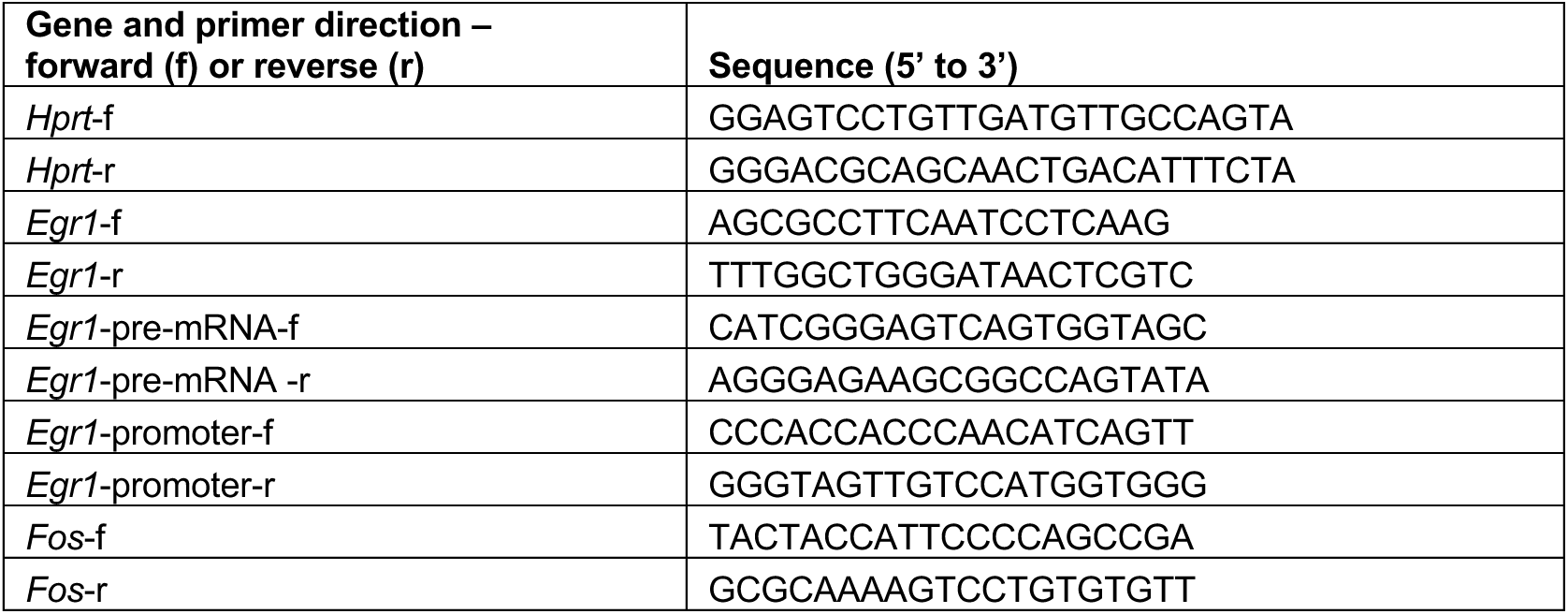
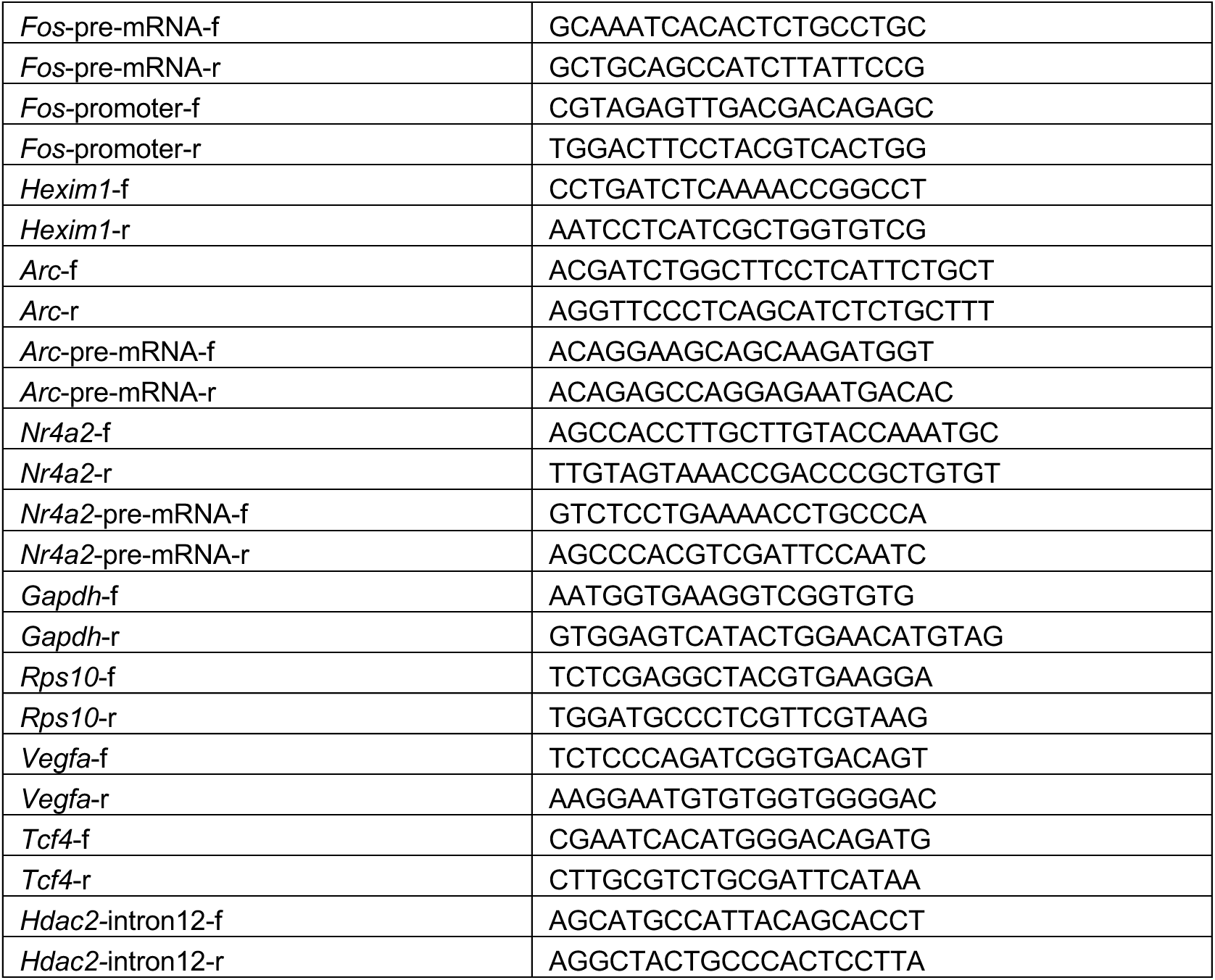

To ensure we only used data from neurons that were healthy enough to robustly activate IEGs, the batch of neurons was only included in the final analysis if *Fos* and *Egr1* mRNAs increased at least 2-fold by KCl in control conditions. This criterion led to the exclusion of eight preparations of neurons across our in vitro studies. Each experiment was run as 2-3 technical replicates, and the average of the technical replicates is reported for each datapoint as an individual biological replicate. For KCl stimulation studies, data was normalized in multiple steps. First, fold changes relative to a housekeeping gene were calculated relative to the plus KCl condition of the control treatment (DMSO, eYFP virus, or water). Then, to preserve variance across experimental conditions, signal from each condition across each biological replicate was summed, then each condition from that biological replicate was divided by that sum. Finally, we calculated the average of the control baseline (minus KCl) condition across biological replicates, and scaled all values by that average. dCas9 experiments were also re-normalized to the non-targeting sgRNA minus KCl condition for each dCas9 condition (NFD, wild-type, or PDND) to account for minor changes elicited by the transfected construct.

#### Immunocytochemistry (ICC)

Cells were plated on PDL-coated cover slips, rinsed twice in ice chilled 1xPBS, and cross-linked in 4% PFA (diluted in 1xPBS) for 15 minutes at room temperature. Cells were blocked and permeabilized in 10% goat serum and 0.3% TritonX-100 in 1xPBS for 1 hr at room temperature. Primary antibodies were added to cells in binding buffer (3.3% goat serum and 0.1% TritonX-100 in 1xPBS). Cells were washed three times in the same buffer used with the primary antibodies. Secondary antibodies were applied in binding buffer for 1 hr at room temperature and washed three times in the same. Cells were washed one time in 1xPBS before mounting in ProLong Gold Antifade mounting solution (Invitrogen). DAPI was either added to the final 1xPBS wash (at 0.5 μg/uL) or supplied in the mounting solution. Slides were dried overnight, images were acquired using a IX73 or Ix81 microscope (Olympus) and cellSens standard software. HEXIM1 antibody was validated for use in ICC with an shRNA. Signal from the HEXIM1 antibody in successfully transfected cells was reduced by about half compared to cells transfected with control shRNAs.

#### Plasmids

eYFP only control and hSyn1:F:Hexim1 expression vectors were cloned into the AAV-shRNA-ctrl backbone (102). AAV-shRNA-ctrl was a gift from Hongjun Song (Addgene plasmid # 85741). The region containing the U6 promoter and shRNA were deleted from constructs. sgRNA were generated as gBLOCKs, PCR amplified, and cloned into the AAV-shRNA-ctrl backbone. pcDNA-dCas9-p300 Core was a gift from Charles Gersbach (Addgene plasmid # 61357 ; http://n2t.net/addgene:61357 ; RRID:Addgene_61357). We changed out the CMV promoter for EF1a. Then, we swapped out the p300 core domain for Hexim1 WT or Hexim1 PDND. Whole plasmid sequencing data was conducted by Plasmidsaurus to validate modifications to these plasmids (**supplementary file:** wholeplasmidsequencingresults_All.txt). dCas9_NFD construct was a gift from John Rinn (Addgene plasmid # 68416 ; http://n2t.net/addgene:68416 ; RRID:Addgene_68416).

#### AAV generation and treatment

Virus was produced using Helper Free Expression System and the 293AAV cell line (Cell Biolabs, Inc.). The cell pellet was lysed by alternating freeze thaw cycles in ethanol/dry ice bath three times. The supernatant was saved, and pellet was additionally extracted with Takara AAV extraction solution. Separately, virus was precipitated from culture media with 10%PEG and 625mM NaCl on ice. The solutions from freeze/thaw, AAV extraction, and the PEG pellet resuspension were combined and purified using iodixanol ultracentrifugation and were concentrated. The viral titer was determined by standard qPCR with the BioRad SYBR Green supermix using serial dilutions of the AAV backbone vector and was concentrated to greater than 1e12 genome copies/mL. 0.3-1uL of virus was applied to cells in 500uL of culture medium 1 week prior to harvesting RNA.

#### Live cell imaging

Cells were imaged with ZOE Florescent Cell Imager (BioRad) 1 week after transduction with AAV.

#### Inhibitors

Inhibitors employed in this study were: DRB (Cayman Chemical, 10010302), FLAVO (Selleckchem, S1230, Batch S123007), JSH-009 (MedChemExpress, Lot 20398), and HMBA (Sigma, lot MKCN8312). DRB applied at dose of 25µM in primary neurons and 50µM in dN2a. FLAVO and HMBA were used at 100nM and 20mM, respectively, in all cell types. Cells were pretreated with DRB, FLAVO, or JSH-009 for 30 min for primary neurons and 30 min to overnight in dN2a. Cells were treated with HMBA for 3 hrs for RT-PCR analysis.

#### Whole cell protein lysate

Cells are washed 1x in PBS, then RIPA buffer supplemented with 1 mM DTT, 1x EDTA-free protease inhibitor (ThermoScientific, Inc.), 200 μM PMSF, 0.1 μM Microcystin LR and 1mM NaF was added to cells, scraped, and collected into a tube. Lysates were incubated on ice for 10 min. with occasional tapping to mix. Samples were centrifuged at 10,000xg for 10 min at 4°C, and the soluble fraction was run in Western blots.

#### Co-Immunoprecipitation

Primary cortical neurons or N2a cells were washed in 1xPBS, then homogenized by scraping in ice cold lysis buffer (150 mM NaCl, 50 mM Tris-HCl, 1% NP-40, 5 mM EDTA, 1 mM DTT, 1x EDTA-free protease inhibitor, 200 μM PMSF, 0.1 μM Microcystin LR and 1mM NaF) and incubated ten minutes on ice. For tissues, two hippocampi were homogenized in the same buffer using a pellet pestle motor and dounce homogenization . The lysates were centrifuged at 12,000xg for 10 min at 4°C. The supernatant was then divided into 3 fractions of 100 μL for IgG, HEXIM1 and CDK9 Co-IPs and 1 fraction of 60 μL was saved for the input sample. An equal mix of protein A and protein G dynabeads (Invitrogen) were washed before use 3 times with Immunoprecipitation (IP) buffer consisting of 150 mM NaCl, 50 mM Tris-HCl, 0.5% NP-40 and 1x EDTA-free protease inhibitor and then we added 1 μL of the individual antibody to 10 μL of the beads. The beads with the antibody and the protein lysate were placed in a tube rotator at 4°C for 1 hr. Bound protein was separated out using a magnetic stand, and a sample of lysate supernatant was saved prior to washing beads. Then, the beads were washed 5 times with IP buffer on the magnetic rack. The proteins were eluted at 65°C for 5 min in agitation with 2x sample buffer (250 mM Tris-HCl pH 8.0, 40% glycerol, 10 mM EDTA, 8% SDS, 400 mM DTT and 0.05% bromophenol blue). These samples were directly loaded into the SDS-PAGE for Western blot analysis.

#### Glycerol Gradient

dN2A cells were treated with 20 mM HMBA for 30 min, then they were harvested and lysed in lysis buffer containing: 150 mM NaCl, 2 mM MgCl_2_, 10 mM HEPES, 5 mM EDTA, 1 mM DTT, 1% PMSF, 1x EDTA-free protease inhibitor, 1 μL/mL RNase inhibitor 40 U/μL, and 0.5% NP-40.

Primary cortical neurons were harvested and lysed in lysis buffer containing either EDTA or calcium. Buffer composition is designed to mimic the intracellular conditions in a resting neuron. Both buffers contained 15 mM NaCl, 2 mM MgCl_2_, 10 mM HEPES, 1 mM DTT, 1% PMSF, 40 μL/mL, 1x EDTA-free protease inhibitor, 1 μL/mL RNase inhibitor 40 U/μL, 0.5% NP-40, 120 mM KCl. In addition to the listed ingredients, EDTA buffer had 5mM EDTA, and calcium buffer had 1 mM CaCl_2_.

After adding the described buffers, the cells were washed in 1xPBS, the protein lysate was placed in a tube rotator for 20 min at 4°C and then centrifuged at 16,000xg for 20 min at 4°C to remove debris. 300 μL of the supernatant were applied to the top of a glycerol gradient. Briefly, the gradient was prepared layering 9 fractions of 500 μL of ascending percentage of glycerol from 10 to 50% (10, 15, 20, 25, 30, 35, 40, 45 and 50%). The column was loaded from the bottom with a 4 cm 21G needle starting with the 10% fraction and going up to 50%. Glycerol fractions were prepared using the same buffer composition (including EDTA or calcium) as its lysis buffer with the exclusion of NP-40 (to avoid foam). The gradients were centrifuged in an Optima L-90K Beckman Coulter ultracentrifuge at 45000 rpm for 16 hours at 4°C. Gradient fractions were stored at -80°C and analyzed by western blot.

To determine the percentage of free and bound forms of each protein, we calculated the summed signal of the protein detected across all fractions and summed the signal in the parts of the gel representing free and bound (as indicated) portions of the gradients. In two EDTA and one Calcium, CDK9 was too faint in the blot to be quantified across the gradients, even though the input had signal. These three experiments were excluded from analysis.

#### Western blots

SDS-PAGE was conducted in 4-20% Mini-PROTEAN TGX Precast Protein Gels (Bio-Rad). Secondary antibodies were goat anti-mouse infrared (IR) 680 (LI-COR Biosciences; #926-68020), goat anti-mouse IR 800 (LI-COR Biosciences; #926-32210), and goat anti-rabbit IR 800 (LI-COR Biosciences; #925-32211). Secondaries used for CoIP were specific to the light chain (Jackson ImmunoResearch) to prevent interference of the antibodies used for the immunoprecipitation when detecting proteins of a similar size to the heavy chain. Membranes were imaged on the LI-COR Biosciences Odyssey fluorescence imaging system, and quantification was done with ImageStudio Software. HEXIM1 and CDK9 antibodies were validated by testing for changes in the intensity of bands of expected sizes using both knockdown shRNAs and recombinant overexpression in N2a and primary neurons. Fold changes in protein expression were calculated such that the control condition normalized to a housekeeping protein equals 100%, then the normalized expression of the treatment are calculated relative to that. To preserve variance across experimental conditions, signal from each condition for each biological replicate was summed, then each condition was divided by the sum. Then, the average of the control condition was scaled to one across the biological replicates.

#### ChIP-qPCR

N2a cells grown and transfected in normal growth media, then swapped to neurobasal complete media 1 day prior to isolation of chromatin. Cells were cross-linked in 1% formaldehyde for 10 minutes at room temperature in buffer containing 10mM HEPES, 10mM NaCl, 1mM EDTA, and 1mM EGTA. Purification of chromatin was done with sequential washing of pelleted material in L1 (50mM HEPES, 140mM NaCl, 1mM EDTA, 1mM EGTA, 0.25% Triton X-100, 0.5% NP40, and 10% glycerol), L2 (10mM Tris-HCl pH8.0, 200mM NaCl), and L3 (10mM Tris-HCl pH8.0, 1mM EDTA, 1mM EGTA) buffers supplemented with 1x EDTA-free protease inhibitor at 4°C. Chromatin was fragmented in L3 buffer plus 10% glycerol using a Qsonica Q800R3 system with 90-100% amplitude for 40-60 minutes on time in cycles of 15s on, 45s off at 4°C. Chromatin was precleared and antibody was prebound with protein A and G Dynabeads. A sample of precleared chromatin was taken for input analysis. Then, remaining precleared chromatin was combined with prebound protein A and G Dynabeads, adding 1x EDTA-free protease inhibitor, 1% DOC, and 1% Triton X-100. Tubes were incubated overnight at 4°C on a rotating stand. Beads were washed twice with low salt (20mM Tris-HCl pH8.0, 2mM EDTA, 150mM NaCl, 1% Triton X-100, 0.1% SDS), once with high salt (20mM Tris-HCl pH8.0, 2mM EDTA, 500mM NaCl, 1% Triton X-100, 0.1% SDS), once with LiCl (20mM Tris-HCl pH8.0, 1mM EDTA, 250mM LiCl, 0.5% Deoxycholate, 1% NP40), and once with TE buffers, removing washes on a magnetic stand. Chromatin was eluted from the beads in TE+1%SDS at 65°C. Crosslinking was reversed overnight at 65°C. Chromatin was purified with proteinase K, extracted with phenol/chloroform, and ethanol precipitated. Resuspended DNA was treated with RNAse A and purified with a PCR purification column. Input samples were diluted to 2.5ng/μL for qPCR, and ChIP samples were diluted to 100μL total volume. Fold enrichment at a site of interest was calculated relative to untreated or NFD-dCas9 input samples using the comparative C_t_ method, normalizing to a primer targeting the 12^th^ intron of *Hdac2* that has little signal in the CDK9 CUT&TAG sequencing dataset.

#### Antibodies

Antibodies.
| Antigen | Company | Source | Dilution | Catalogue | Lot | Isotype |
| --- | --- | --- | --- | --- | --- | --- |
| HEXIM1 | abcam | Rabbit | 1:1000 (WB/ICC)<br>6μg/IP (ChIP) | ab25388<br>1mg/mL | GR3245992-1 or<br>GR3460630-1 | IgG |
| Beta-ACTIN | Cell Signaling | Mouse | 1:2000 (WB) | 8H10D10 | 21 | IgG2b |
| CCNT1 | Cell Signaling | Rabbit | 1:1000 (WB)<br>3μg/IP (ChIP) | 81464<br>D1B6G | 1 | IgG |
| CDK9 | abcam | Rabbit<br>monoclonal | 1:1000 (WB)<br>3μg/IP (ChIP) | ab239364 | 1023428-1 | IgG |
| FLAG (M2) | Sigma | Mouse | 1:1000 (WB) | F1804 |  | IgG1 |
| Tubulin,<br>beta III<br>(Tuj1) | Millipore | mouse | 1:2000 (ICC) | TU-20,<br>MAB1637 | 3407183 | IgG1 |
| NeuN | Abcam | Mouse | 1:500 | ab104224 |  | IgG2b |
| GAPDH | Abcam | Mouse<br>monoclonal | 1:2000 (WB) | ab8245 | 1035914-4 | IgG1 |
| Rabbit IgG | Thermofisher | Rabbit | NA | 02-6102 | SJ257848 | IgG |
| RNAP2-pS2 (3E10) | Millipore<br>Sigma | Rat | 1:500 (WB) | 04-1571 |  | IgG1 |
| RNAP2-pS5 (3E8) | Millipore<br>Sigma | Rat | 1:500 (WB) | 04-1572 |  | IgG2a |
| Total<br>RNAP2 (F-12) | Santa Cruz<br>Biotechnology | Mouse | 1:200 (WB) | sc-55492 |  | IgG2b |

#### Statistics

RT-PCR and blot quantitation data was analyzed statistically in GraphPad Prism Version 10.

### Data availability

ROSMAP resources can be requested at https://www.radc.rush.edu and www.synpase.org.

## Supporting information

Plasmid Sequencing

Supplementary Table 1. Demographic data for the ROSMAP bulk RNAseq

## Acknowledgements

We thank Hero Haji for her help with cloning and Eugenia V. Gurevich for help with virus generation.

## Author contributions

**Myo Htet**: Investigation; Methodology; Resources; Visualization; Writing – review & editing. **Camila Estay-Olmos**: Investigation; Methodology; Visualization; Writing – review & editing. **Lan Hu**: Investigation; Resources. **Yiyang Wu**: Formal analysis; Visualization. **Brian E. Powers**: Investigation. **Clorissa D. Campbell**: Investigation. **Aishwarya Rameshwar:** Investigation. **Mohamed R. Ahmed**: Methodology; Resources. **Timothy J. Hohman**: Conceptualization; Funding acquisition. **Yanling Wang**: Data curation; Formal analysis. **Julie A. Schneider**: Resources. **David A. Bennett**: Data curation; Funding acquisition; Resources; Writing – review & editing. **Vilas Menon**: Resources. **Philip L. De Jager**: Data curation; Formal Analysis. **Garrett A. Kaas**: Methodology; Supervision. **Roger J. Colbran**: Funding acquisition; Methodology; Supervision; Writing – review & editing. **Celeste B. Greer**: Conceptualization; Formal analysis: Funding acquisition; Investigation; Methodology; Resources; Supervision; Visualization; Writing – original draft; Writing – review & editing.

## Funding and additional information

This study is supported by NIH grants K01 MH129760 to CBG, R01 MH057014 to RJC, T32 MH065215 to RJC, R01 AG074012 to TJH, R01 AG061518 to TJH, and R01 AG059716 to TJH. ROSMAP is supported by P30 AG10161, P30 AG72975, R01 AG15819, R01 AG17917, U01 AG46152, and U01 AG61356. The content is solely the responsibility of the authors and does not necessarily represent the official views of the National Institutes of Health.

