## Supplemental Figures for "HEXIM1/P-TEFb complex controls RNA polymerase II pause release and immediate early gene induction following neuronal depolarization"

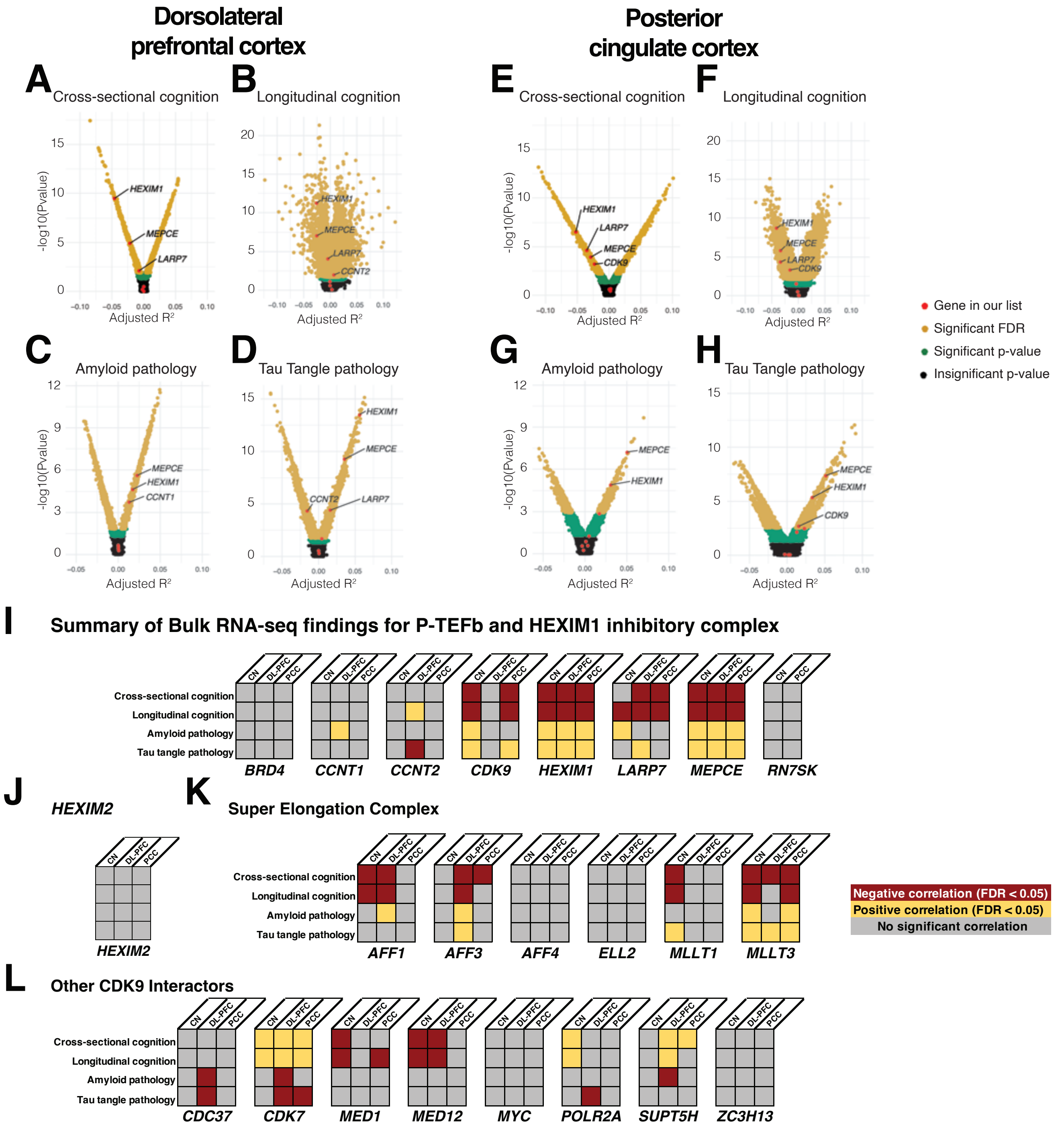


**Supplemental Figure 1. Correlations between P-TEFb regulatory complex components and AD-related phenotypes.** Correlations of *BRD4*, *CCNT1*, *CCNT2*, *CDK9*, *HEXIM1*, *LARP7*, and *MEPCE* mRNAs with (**A**) cross-sectional cognition, (**B**) longitudinal cognition, (**C**) amyloid pathology, and (**D**) Tau tangle pathology in dorsolateral prefrontal cortex. Correlations of *BRD4*, *CCNT1*, *CCNT2*, *CDK9*, *HEXIM1*, *LARP7*, and *MEPCE* mRNAs and *RN7SK* non-coding RNA with (**E**) cross-sectional cognition, (**F**) longitudinal cognition, (**G**) amyloid pathology, and (**H**) tau tangle pathology in posterior cingulate cortex. Individual dots represent a single gene, and genes in the yellow region of the volcano plots are significantly correlated with the indicated pathology (FDR < 0.05). (**I**) Summary of correlations in gene list with FDR < 0.05 in bulk sequencing datasets across three brain regions. (**J-L**) Summary of correlations in (**J**) *HEXIM2*, (**K**) super elongation complex components, and (**L**) other CDK9 interactors (identified with STRING database (1)) with FDR < 0.05 in bulk sequencing datasets across three brain regions.

1. Szklarczyk, D., Kirsch, R., Koutrouli, M., Nastou, K., Mehryary, F., Hachilif, R. *et al.* (2023) The STRING database in 2023: protein-protein association networks and functional enrichment analyses for any sequenced genome of interest Nucleic Acids Res **51**, D638-D646 10.1093/nar/gkac1000

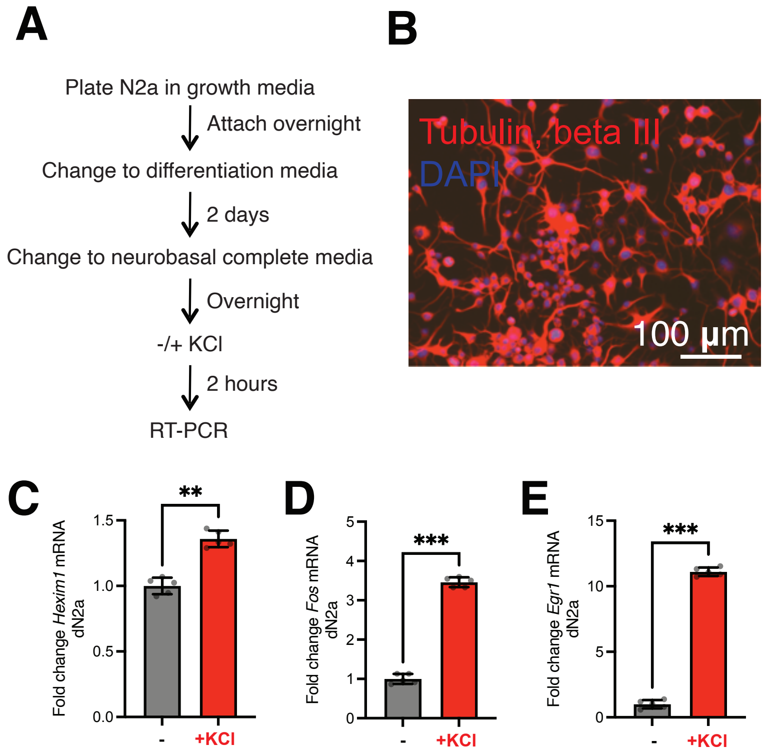


**Supplemental Figure 2. N2a cell differentiation procedure and effect of KCl stimulation on *Hexim1* mRNA and IEGs.** (**A**) Timeline of N2a plating, differentiation, and stimulation. (**B**) ICC of a neuronal marker (TUJ1) to show cell morphology in dN2a. (**C-E**) Fold changes in mRNA expression calculated relative to *Hprt* mRNA (housekeeping gene) following KCl stimulation in (**C**) *Hexim1* (**D**) *Fos* and (**E**) *Egr1* mRNA after KCl stimulation in dN2a (RT-PCR). n=5 biological replicates. Paired two-tailed t test. ** $p$<0.01. Error bars represent standard deviation.

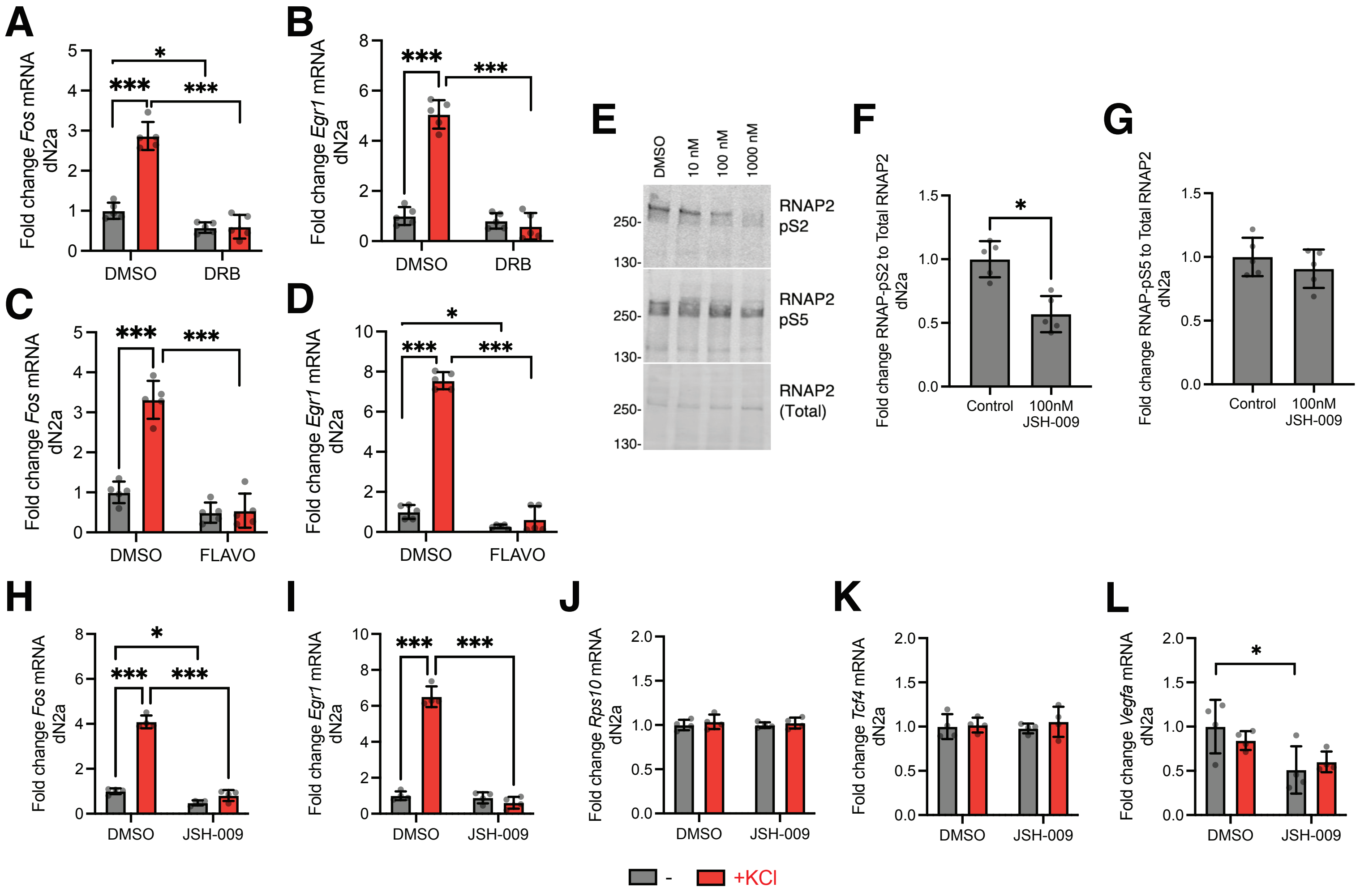


**Supplemental Figure 3. P-TEFb regulates activity-dependent IEG induction in dN2a.** (**A**) *Fos* and (**B**) *Egr1* mRNA levels (RT-PCR) after pretreatment with DRB followed by KCl stimulation in dN2a. n=5 biological replicates. (**C**) *Fos* and (**D**) *Egr1* mRNA levels (RT-PCR) after pretreatment with FLAVO followed by KCl stimulation in dN2a. n=5 biological replicates. (**E**) Effect of 2.5 hr treatment of dN2a with indicated doses of JSH-009 on RNAP2 phosphorylation at serine 2 (pS2) and serine 5 (pS5). Note that phosphorylation increases the apparent molecular weight of RNAP2, so Total RNAP2 mainly shows the unphosphorylated form (just above 250kDa), and the phospho-specific antibodies detect slightly higher molecular weight bands. (**F**) Quantitation of RNAP2-pS2 and (**G**) RNAP2-pS5 relative to Total RNAP2 following 2.5 hr. 100nM JSH-009 treatment. n=5 biological replicates. Paired Student's t test. $^*p<0.05$. (**H**) *Fos,* (**I**) *Egr1,* (**J**) *Rps10*, (**K**) *Tcf4*, and (**L**) *Vegfa* mRNA levels (RT-PCR) after pretreatment with 100nM JSH-009 followed by KCl stimulation in dN2a. n=4 biological replicates. All RT-PCR data normalized to *Hprt* housekeeping gene. Two-way ANOVA with Sidak's multiple comparisons test. $^*p<0.05$, $^{***}p<0.001$. Error bars represent standard deviation.

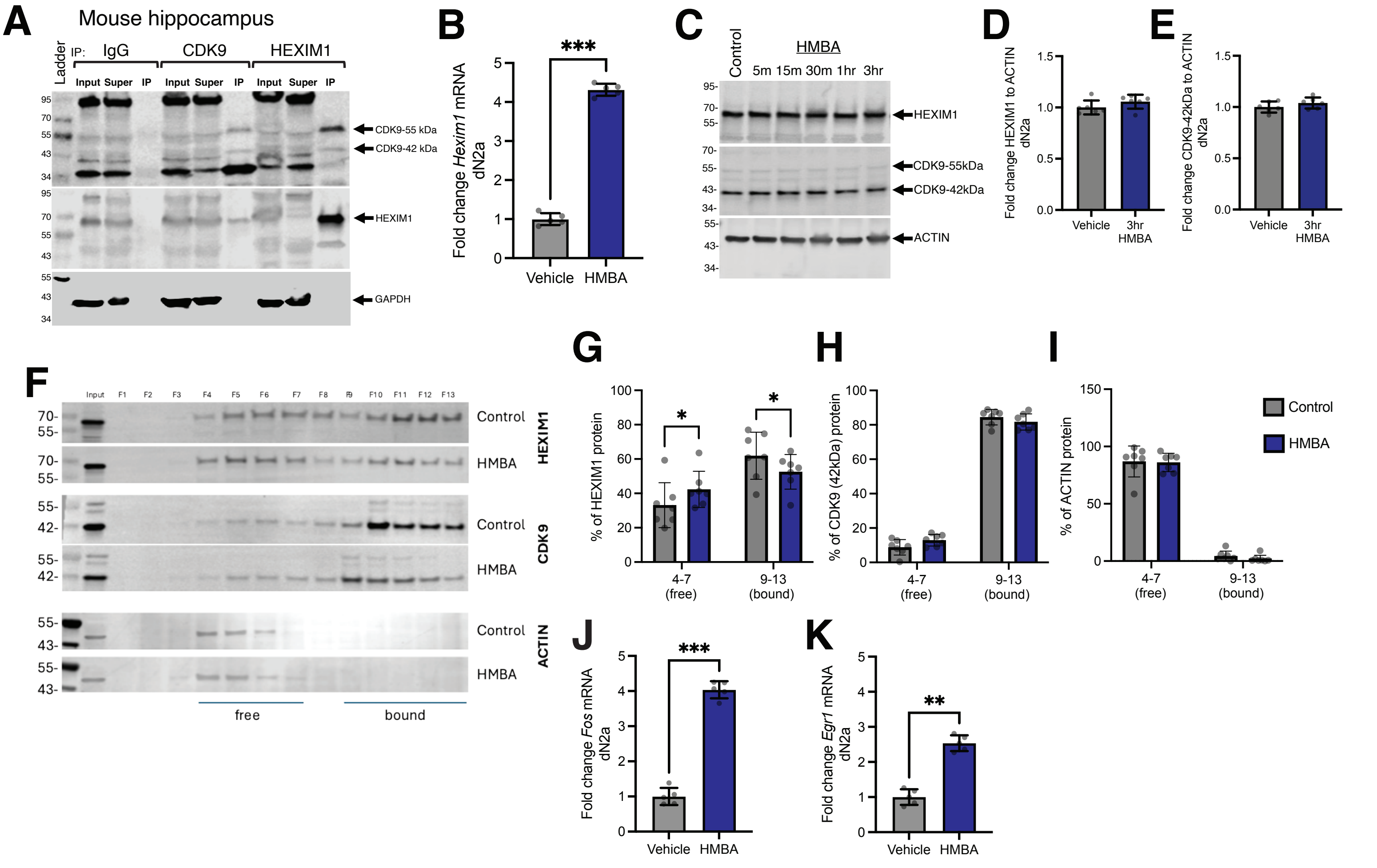


**Supplemental Figure 4. P-TEFb associates with HEXIM1 in vivo, and HMBA affects baseline IEG expression in dN2a.** (**A**) Coimmunoprecipitations using IgG, CDK9, and HEXIM1 antibodies from mouse hippocampus protein lysate. Input is the lysate prior to immunoprecipitation, Super = supernatant from the immunoprecipitation, and IP = immunoprecipitated material after bead washing. (**B**) *Hexim1* mRNA expression changes after HMBA treatment (RT-PCR). n=5 biological replicates. Paired Student's t test. (**C**) HEXIM1, CDK9, and ACTIN protein changes at indicated time points up to 3hr. after the start of HMBA treatment (Western blot). (**D-E**) Quantitation of (**D**) HEXIM1 or (**E**) CDK9 (42kDa) levels relative to ACTIN at 3hr timepoint (n=6 biological replicates). Not significantly different by paired Student's t test ($p > 0.05$). (**F**) Glycerol gradient of protein lysates following HMBA treatment. (**G-I**) Quantitation of relative fraction of (**G**) HEXIM1, (**H**) CDK9 (42kDa) and (**I**) ACTIN protein in 'free' low-molecular weight fractions (4-7) and 'bound' high molecular weight fractions (9-13). n = 7 biological replicates. Two-way ANOVA with Sidak's post hoc. (**J-K**) mRNA expression changes relative to *Hprt* in dN2a after HMBA treatment (RT-PCR) for (**J**) *Fos* and (**K**) *Egr1*. n=5 biological replicates. Paired Student's t test *$p$<0.05, **$p$<0.01, ***$p$<0.001. Error bars represent standard deviation.

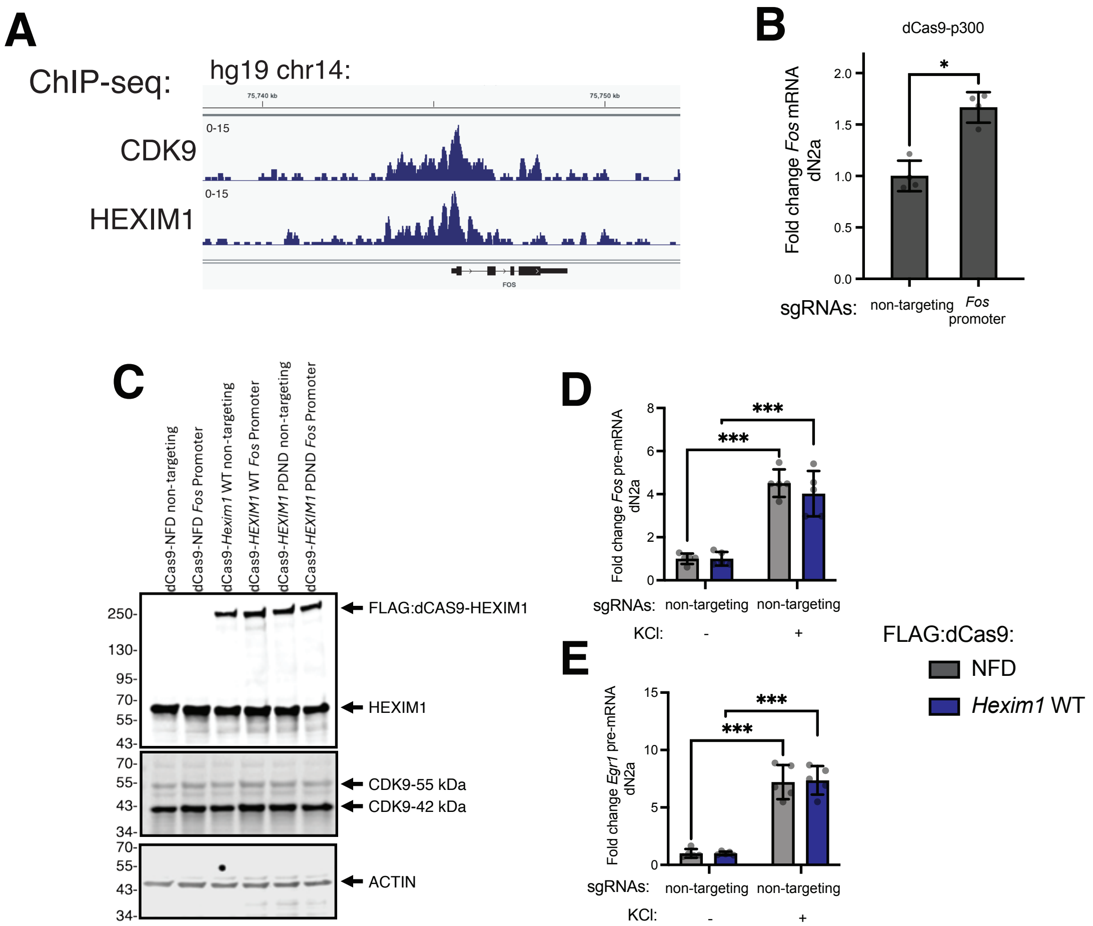


**Supplemental Figure 5. CDK9 and HEXIM1 are bound to *Fos* promoter, and dCas9 vector expression.** (**A**) ChIP-seq track reads from A375 cells. Data from a previously published CDK9 and HEXIM1 datasets around *FOS* gene in the human genome (2). Reads at the *FOS* gene were visualized with Integrated Genome Viewer. (**B**) Test of sgRNA targeting to the *Fos* promoter using dCas9-p300 transcriptional activator in N2a cells. *Fos* mRNA measured by RT-PCR and normalized to *Hprt* housekeeping gene. Paired t test. n=4 biological replicates. **p*<0.05. (**C**) Western blots of transfected N2a to test expression of the HEXIM1 containing constructs relative to endogenous HEXIM1. Protein lysates were blotted with antibodies against HEXIM1 (top), CDK9 (middle), and ACTIN (bottom). (**D**-**E**) Effect of transfecting dCas9-HEXIM1 WT without targeting it to a specific genomic locus on (**D**) *Fos* and (**E**) *Egr1* expression. Fold change is identified relative to *Hprt* housekeeping gene. Two-way ANOVA with Sidak's multiple comparisons test. n=5 biological replicates. ****p*<0.001. Error bars represent standard deviation.

2. Tan, J. L., Fogley, R. D., Flynn, R. A., Ablain, J., Yang, S., Saint-Andre, V. *et al.* (2016) Stress from Nucleotide Depletion Activates the Transcriptional Regulator HEXIM1 to Suppress Melanoma Mol Cell **62**, 34-46 10.1016/j.molcel.2016.03.013

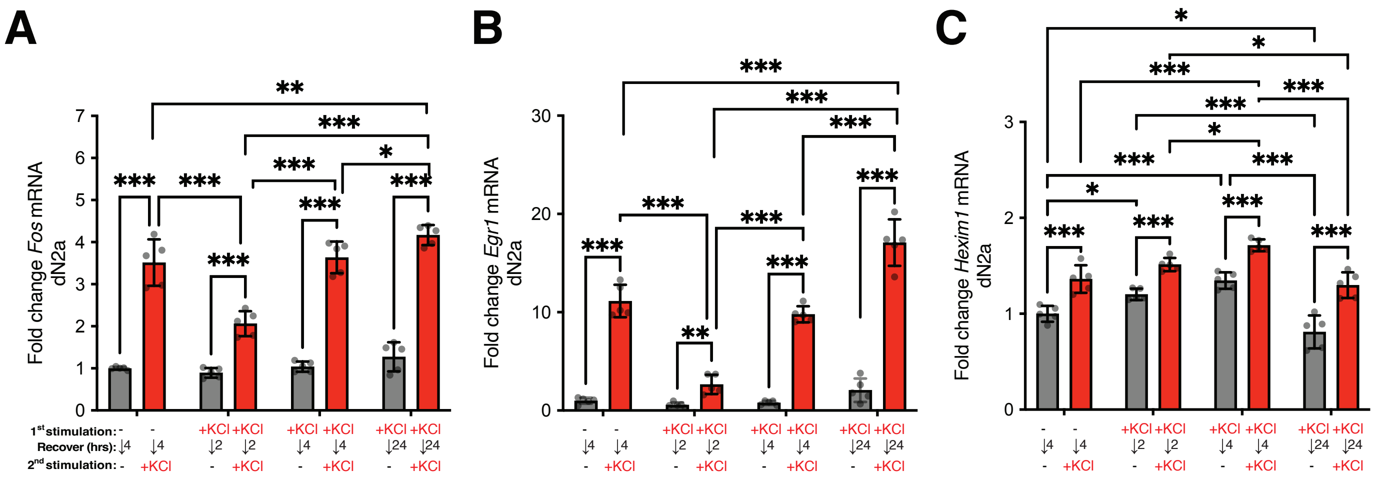


**Supplemental Figure 6. Transcriptional responses of IEGs are dampened following a prior stimulation in dN2a, whereas *Hexim1* mRNA induction increases.** (**A-C**) IEG responses of (**A**) *Fos*, (**B**) *Egr1*, and (**C**) *Hexim1* mRNA (RT-PCR; fold change relative to *Hprt*) in dN2a treated with 50mM KCl according to timeline described in **Fig. 7M**. n=5 biological replicates. Two-way ANOVA with Tukey multiple comparisons test. n=5 biological replicates. *$p$<0.05, **$p$<0.01, ***$p$<0.001. Error bars represent standard deviation.

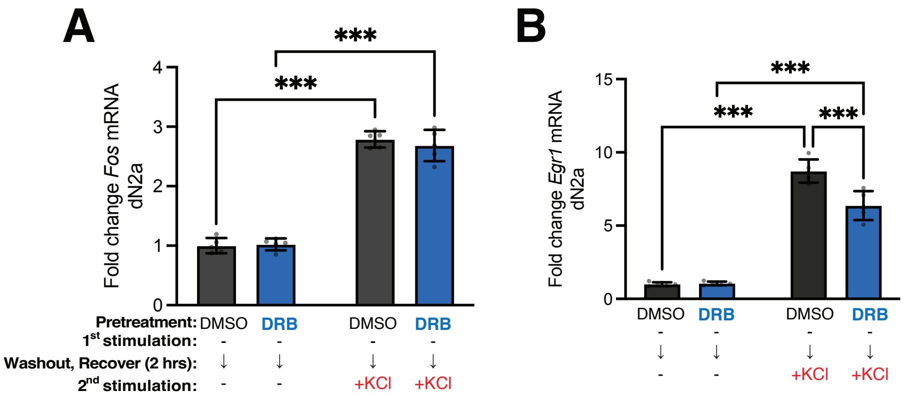


**Supplemental Figure 7. Effect of DRB pretreatment then washout on *Fos* and *Egr1* inducibility in dN2a.** (**A-B**) Following the timeline depicted in **Fig. 8A**, we also measured the effect of vehicle (DMSO) or DRB pretreatment on (**A**) *Fos* and (**B**) *Egr1* mRNA expression levels (RT-PCR; fold change relative to *Hprt*) in unstimulated (left two bars) or stimulated (right two bars) dN2a. Calculated as a fold change relative to KCl treatment during the second stimulation with no prior drug or KCl treatments. Two-way ANOVA with Sidak's multiple comparisons test. n=5 biological replicates. ***$p$<0.001. Error bars represent standard deviation.
